# Transition from infectivity and immune escape to pure escape as an evolutionary strategy during the COVID-19 pandemic

**DOI:** 10.64898/2026.02.26.706090

**Authors:** Benjamin Kotzen, Sarah Gurev, Noor Youssef, Javier Jaimes, Jeremy Luban, Debora Marks, Michael Seaman, Jacob E. Lemieux

## Abstract

New SARS-CoV-2 variants have undergone repeated selective sweeps since the beginning of the COVID-19 pandemic, but the fitness advantages and mechanisms driving these sweeps are not fully understood. We developed a probabilistic modeling framework to analyze pandemic growth, infectivity, and immune escape, explicitly accounting for seven immune exposure histories in 5,732 experiments and estimating the effects of 835 mutations. We found infectivity was important for early variants, but as gains became zero-sum, growth became driven by consistent increase in immune escape conferred by a primarily additive effect of mutational accumulation. While phenotypic tradeoffs exist for individual mutations, successful viral strains boast assemblages of mutations that do not sacrifice infectivity for escape. Thus, during an apparent transition to endemicity, SARS-CoV-2 evolution ascended along an evolutionary ridge in the mutational space defined by infectivity and escape, with infectivity reaching an early peak and antigenicity continuing to evolve.

## INTRODUCTION

The COVID-19 pandemic, caused by the virus SARS-CoV-2, has been a cause of death and serious illness since its emergence in late 2019. The virus has been able to spread due to constant mutation, leading to the emergence of genetically and antigenically divergent variants. Effective pandemic preparedness for SARS-CoV-2 relies on understanding and predicting the ability of novel strains to escape host immune responses(1–5), as immune escape has ramifications for vaccine development, course of infection, and transmission dynamics.

Many methods exist to examine the effects of mutations on immune escape, including Deep Mutational Scanning (DMS)(6), neutralization assays(5), and AI-powered evolutionary escape modeling(1). In DMS, a library of genomes is synthesized such that every possible amino acid mutation is represented and measured for its impact on receptor binding(3,7,8), surface expression(3,7,9), or antibody binding(2–4,8,10–26). Computational models exist that assess mutations at a scale not possible with current experimental approaches. For example, EVEscape incorporates fitness measures by training a variational autoencoder on homologous Spike proteins across coronaviruses along with biochemical and structural measures relevant to immune escape. Other approaches, such as BVAS and PyR0 rely on pandemic sequencing to measure relative growth rates of viral strains and infer mutational fitness. In this work, we present another high-throughout computational approach that fills in the gaps of these existing methods that can interpret diverse, non-homogenized experimental neutralization assays at scale to identify key single mutation effects.

We focus our investigation on the Spike protein of SARS-CoV-2, which is the primary modulator of interaction with the host environment, including antibodies and host receptors(27). The 1273-amino-acid-long Spike protein includes several subdomains such as the N-Terminal Domain (NTD, amino acid positions 13-304), Receptor Binding Domain (RBD, amino acid positions 319-541), Receptor Binding Motif (RBM) within the RBD (amino acid positions 438-508), S1/S2 cleavage region (amino acid positions 672-709), Fusion Peptide (FP, amino acid positions 788-806), Internal Fusion Peptide (IFP, amino acid positions 816-733), Heptad Repeat 1 (HR1, amino acid positions 918-983), and Heptad Repeat 2 (HR2, amino acid positions 1,162-1,203)(28). It has been shown by previous research that nearly all anti-SARS-CoV-2 antibodies target the RBD or NTD(29). Antibodies targeting the RBD have been found to make up 90% of neutralizing activity in COVID-19 serum(11,30,31). The NTD, particularly the NTD supersite, is also a strong target for antibodies(29,30,32).

To disentangle the impact of mutations on fitness or antibody escape for SARS-CoV-2, we generated the first mutation-level immune escape dataset using hierarchical Bayesian regression that decomposes single mutation effects from diverse, multi-mutant neutralization data. We show that there are relevant differences across humoral immune responses which have the potential to become increasingly disparate as the illness becomes more epidemic, shifting to a seasonal virus. These results have implications for vaccine development, forecasting viral growth and public health policy. Our approach uniquely allows us to inspect model parameters, including those typically seen as “nuisance parameters” for both interpretability and biological insights.

## RESULTS

### Quantifying mutation-specific contributions to immune escape

To quantify immune escape at the mutation-level, we analyzed our previously published neutralization data (5) and data aggregated and made public in the Stanford Coronavirus Antiviral & Resistance Database by Tzou et. al., 2020 (33). In sum, we analyzed 5,223 Spike proteins including 779 total amino acid mutations relative to B.1 (D614G), arranged in 383 unique sequences. We built a Bayesian hierarchical linear regression model to describe the level of polyclonal antibody escape, measured as the log_10_-fold-reduction in NT50 neutralizing titer (Figure 1A-B).

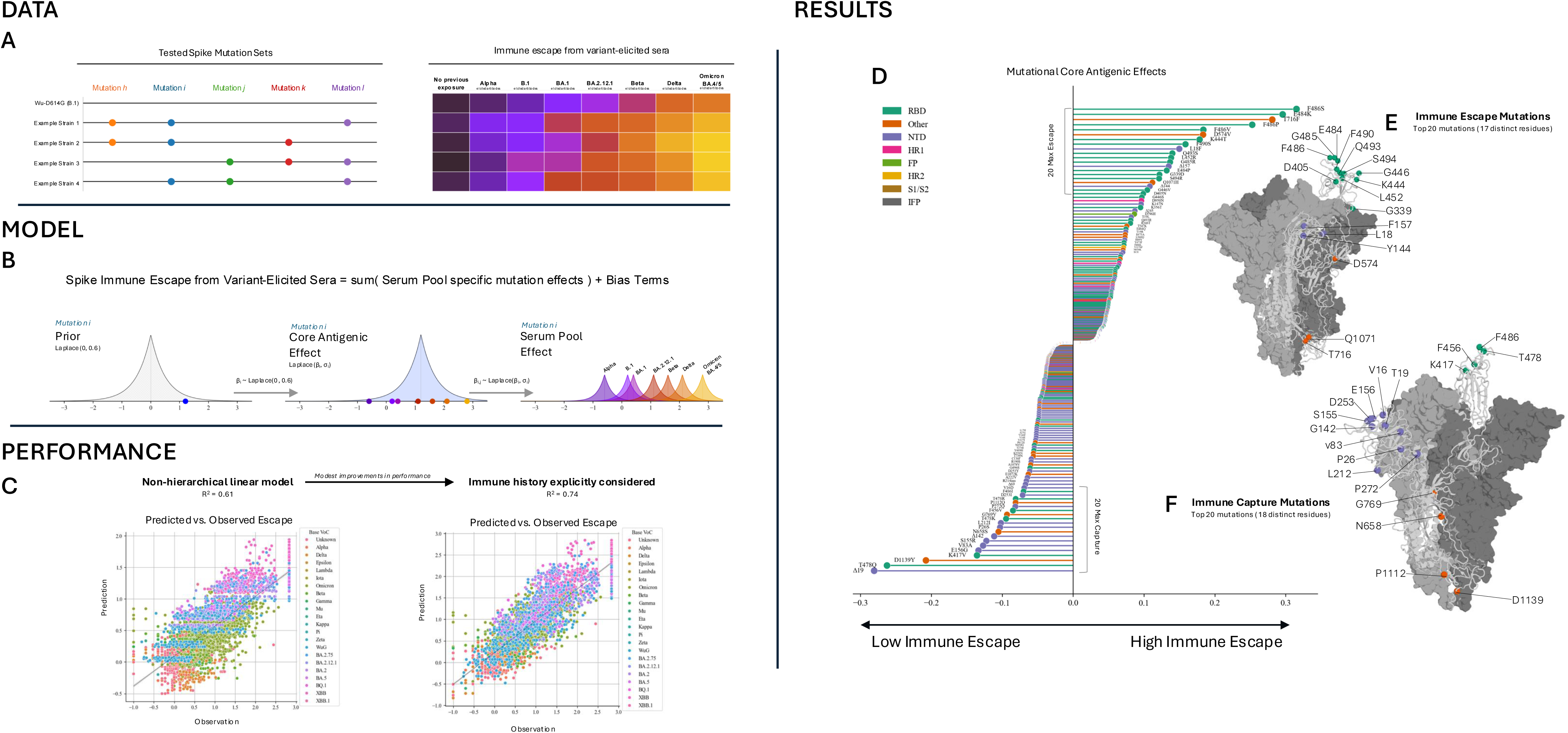

Mutations associated with the greatest escape were concentrated densely in the RBD (Figure 1D-E, Supplemental Figure S1). Of the ten mutations with the greatest effect, seven are in the RBD. Notably, three are at residue F486, a key residue for ACE-2 binding (11,15,34), one of the most exposed residues of the entire protein and a target of many antibodies (Figure 1E, Supplemental Figure S2) (12,14,15,18,19,21,22,34). One of the top ten mutations, L18F, is in the NTD, and two of the top-ten mutations, T716F and D574V, are outside both these regions.

While some high-escape mutations outside the NTD and RBD such as D405N (and G339D to a lesser extent) are still moderately accessible to antibodies, the mutations T716F, D574V, and Q1071H contribute to high immune escape despite being farther down the stalk of the Spike protein where they are less accessible to antibodies. Mutations in the NTD contributed to immune escape, especially the deletions at residues Y144 and F157, plus the substitution L18F. We analyzed the weighted contact number (WCN) as a measure of accessibility, and the binding sites of nearly 300 known antibodies (1) and found that the set of mutations that confidently escaped immune responses (95% CI of effect greater than 0) were more exposed (p<0.05) and targeted by more antibodies (p<0.05) than mutations that did not confidently contribute to escape / capture (mutations with 95% CI of effect including 0).

However, the NTD is also densely populated with mutations leading to immune capture, i.e. mutations that lead to better neutralization by improving antibody binding. Of the 10 most easily captured mutations, six are in the NTD: Δ19, E156G, V83A, S155R, Δ142, P26S. (Figure 1D,F); we observe particular density of immune-capture mutations, especially deletions, in the N3 loop / supersite beta-hairpin of the NTD (residues 140-158), which has previously been shown to be a common SARS-CoV-2 epitope (30). Certain RBD mutations also attenuate immune escape, such as T478Q and K417V, as well as other mutations outside the RBD, for example D1139Y, P1112Q, and N658S. Parsing through data from GISAID (35), we found that all these mutations were seen early in the pandemic (pre-August 2020), allowing many antibodies to develop targeting these mutations. Similar to T716F, which aided in immune escape, D1139Y and P1112Q are on the stalk of the Spike protein, indicating that they are inaccessible to antibodies directly but may alter the conformation of the protein (Figure 1E). We analyzed surface exposure (WCN) and antibody binding activity at the residues of mutations we are confident are captured by immune responses (mutations with 95% CI less than 0) and found that these residues are more likely to be exposed (p<0.05) and targeted by antibodies (p<0.05) than mutations with effect CIs that include 0.

In our Bayesian regression model, we applied intercept terms to correct for experimental or serological conditions likely to influence measured escape values, many of which have an impact on the same order of magnitude as an individual mutation with a large contribution (Supplemental Figure S3). We found neutralization titer to be improved by having convalescent plasma or having more than two previous exposures to SARS-CoV-2 virus. Unexpectedly, having a most recent exposure more than six months prior to serum isolation corresponded to an increase titers, though this is likely confounded by undocumented secondary SARS-CoV-2 infection after vaccination, as titers are known to be higher at 1 or 2 months than 6 months(36). Assay type also influenced measured escape, with greater immune escape measured in pseudovirus and hiVNT assays, and lower escape measured in VSV chimeric virus or SARS-CoV-2 recombinant assays.

While the full hierarchical bayesian model reveals relevant biology and shows strong correlation between prediction and observation (Figure 1C), we note that a simplification of the model to use only the core antigenic effect (ignore hierarchical layers accounting for immune history) still performs well, yielding a correlation between prediction and observation with R^2^=0.61.

### Identifying specific epitopes targeted by different infection histories

Our hierarchical model assumed the core antigenic effect for each mutation was at the center of a cluster of several fine-tuned effects which reflect variation in immune responses across polyclonal sera with different immune histories (Figure 2: Illustration). In fitting the model, we pooled the data to define seven immune history pools based on the variant of most recent exposure: Alpha, B.1, BA.1, BA.2.12.1, Beta, Delta, and Omicron BA.4/5.

**Figure 2:**
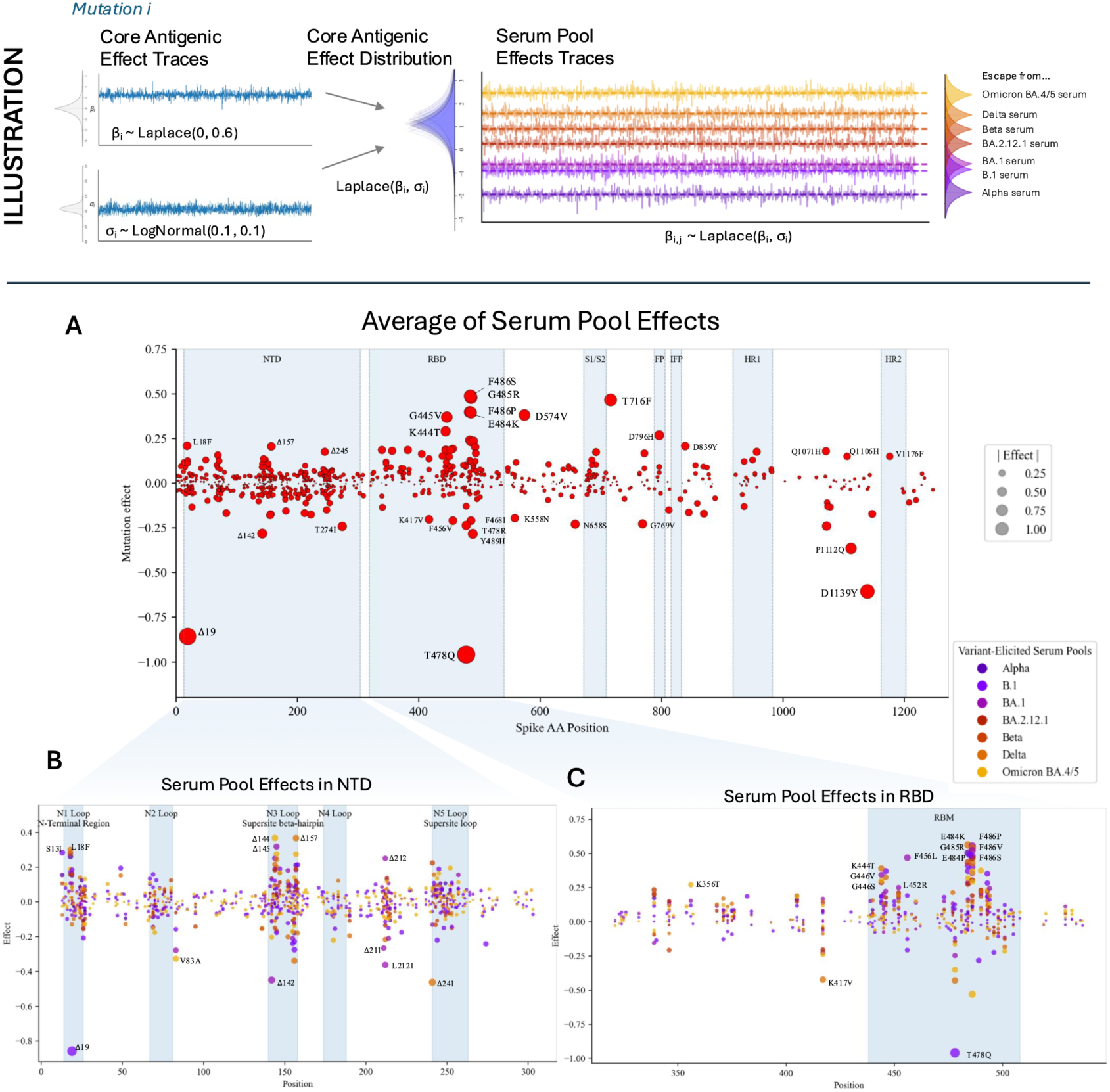
Analysis of serum pool effects. (ILLUSTRATION) For each mutation, a core antigenic mean effect and standard deviation are sampled; the resulting Laplace distribution characterized by these parameters is the prior distribution from which serum pool effects are then sampled. (A) Most immune activity occurs in the NTD and RBD, including high immune capture of the deletion at residue 19 or the substitution T478Q; there is still significant antigenic activity outside these regions as well. (B) Mutations in N1, N3, and N5 “antigenic supersites” often evade immune responses from B.1-exposed individuals; mutations in N3 and between N4 and N5 (around residues 211-222) continue to show antigenic activity in other serum pools. (C) Effects of specific substitutions within the RBD are modulated by serum pool / immune history. Sometimes, different substitutions to the same site produce opposite serum pool specific effects (e.g., K417N versus K417T, K417V, K417R).

We investigated these serum pool-specific coefficients to quantify the extent to which immune history leads to differences in mutational effects. In general, several mutations to the RBD escaped immune reactions elicited by all previous exposures, and NTD substitutions / deletions were often targeted by specific serum pools (Figures 2A-C). We found that many of the most extreme cases of immune escape / capture occurred for the B.1 (S:D614G)-elicited serum pool, especially in the NTD and RBD (Figures 2B-C, Supplemental Figure S4). Within the NTD, most of the escape activity from this serum pool was concentrated in the N1, N3, and N5 supersites, as has been previously reported(29,30,32). We also uncovered immune activity outside these regions, notably with the mutation V83A right outside the N2 loop which leads to immune capture in the Omicron BA.4/5, BA.1, and B.1 serum pools. Various substitutions and deletions between the N4 and N5 loops (specifically sites 211-212) were found to show strong immune escape / capture phenotypes, especially in the BA.1 serum pool (Figure 2B). In the RBD, several mutations exhibited immune escape or capture modulated by the immune history of the serum pool (Figure 2C). The RBD (especially the RBM) contained many mutations that were specifically targeted by BA.4/5-elicited responses, such as F486I and T478R. Generally, there seemed to be better capture of NTD mutations by serum elicited from B.1 and BA.1, and there was better capture of RBD mutations by serum elicited from Delta and Omicron BA.4/5.

The B.1 (D614G)-elicited response uniquely captures Δ19 and T478Q, as well as the pair of mutations D1139Y and P1112Q which appear at the base of the spike stalk (Supplemental Figure S4). The B.1.1.7 (Alpha variant)- and B.1.351 (Beta variant)-elicited responses are uniquely well evaded by T716F, E484K, and L18F. We summarized the patterns in these immune responses by performing PCA on the serum-pool-level mutation effects (Supplemental Figure S5). The result of this analysis further demonstrates the similarity between B.1.1.7 (Alpha)- and B.1.351 (Beta)-elicited immune responses and shows that the greatest differences in immune responses are between B.1 (D614G), B.1.617.2 (Delta variant), and B.1.1.529.4/5 (Omicron BA.4/5 variants).

To further probe the differential responses across these immune histories, we also described the difference in escape of a mutation between two immune responses as a random variable (Supplemental Figure S6: Top 8 Mutations/Pools with Pool-Specific Effect Differentials). This approach quantifies the evidence of a difference in escape between the two responses. We found that most of the mutations with evidence of differential responses across immune histories were located in the RBD, NTD, and HR2, indicating that RBD and NTD are key epitopes for antibodies and that mutations in the HR2 can alter antibody binding (Figures 2B-C, Supplemental Figure S6, Supplemental Figure S7). Deletions in the NTD had statistically more disparate effects than substitutions in the NTD (independent t-test, p<0.05), while deletions elsewhere did not show a statistically significant difference across immune histories compared to substitutions. We found the mutations with the highest probability of having a true difference in escape across two immune responses were often at residues K417, T478, and F486 (Supplemental Figure S6). K471V escapes B.1 (S:D614G) but is captured by B.1.617.2 (Delta) and B.1.1.529.4/5 (Omicron BA.4/5); the same pattern holds for T478K. F486I escapes B.1 but is captured by B.1.1.529.4/5.

### Variance estimates detect nonlinear interactions among mutations

While our modeling approach explicitly accounts for variability in the effects of mutations across immune histories, the variability in the effects of mutations due to nonlinear interaction with other mutations is captured implicitly by the variance in the parameter estimates for each mutation *i* (var(β_i_)) (Figure 3A). Because this variance can come from many sources (variance in presence/absence of mutation *i*, var(X_i_); size of dataset, n; colinearity, R_i_^2^; or true variance in effect, s_i_^2^), we used variance decomposition (Figure 3A)(37) to estimate s_i_^2^ for each mutation *i*, the amount of true variance in the core antigenic effects of mutations based on their interactions with other mutations (Figure 3B). The resulting set of high-variance mutations is different from mutations highlighted by other measures in a way that is both statistically and biologically significant (Figure 3B). Mutations such as R346T and L452R had relatively stable measured effects, but when variation in that effect was decomposed, it was found that these mutations likely have strong variation depending on their context (Supplemental Figure S8). Other mutations, such as G261F, had high measured variance that was likely due to being seen rarely in the data (G261F was only seen four times); by decomposing the variance measurement, the estimated true variance of mutations like this was greatly reduced. As a result, most mutations were found to have low true variance in core antigenic effect after adjusting measured variance. There is a manifold of mutations where measured variance correlates well with true variance (Supplemental Figure S8).

**Figure 3:**
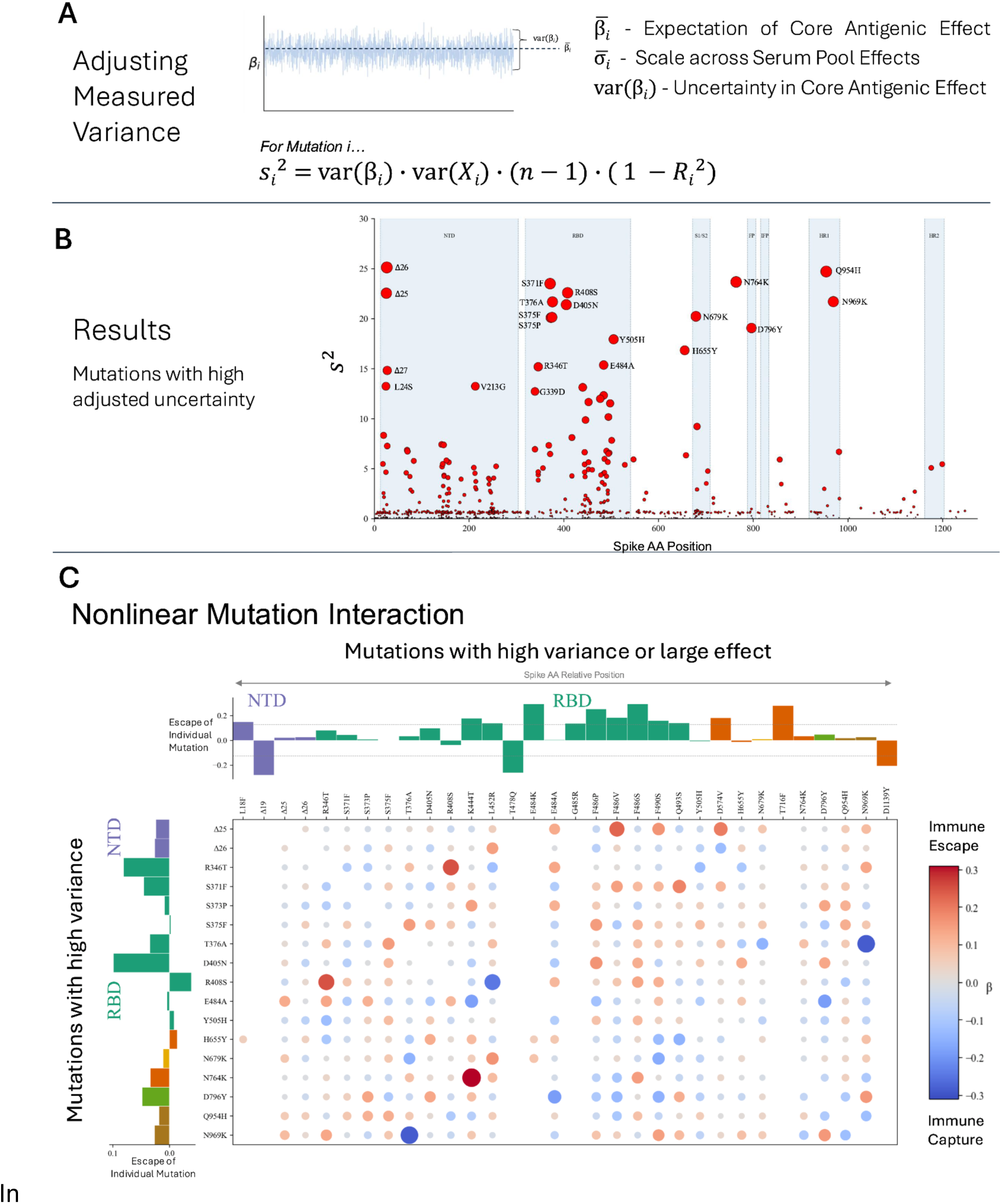
Variance decomposition and nonlinear interactions among mutations. (A) Variance in core antigenic effect, measured directly from the MCMC trace, can be decomposed using (37) to yield an estimated true variance in effect that accounts for factors that may artificially inflate or reduce variance measurements. (B) Mutations with the highest adjusted variance in effect are often in the low-binding region of the RBD (outside the RBM). (C) Nonlinear pairwise mutation interactions may be strong even when the individual mutation effects are weak or the mutation pair spans multiple regions of the protein.

We compared these estimates of true variance with DMS measurements of immune escape of mutations on different background strains. We aggregated 20 DMS studies measuring immune escape(2–4,8,10–25), normalizing escape measurements per study, antibody, and background variant. We then took the maximum normalized escape value across studies and antibodies to get a representative escape value per mutation per background strain. The variance in these values across background strains was somewhat correlated (R^2^ = 0.31) with estimated true variance using variance decomposition (p<0.05) (Supplemental Figure S9).

While mutations with large effects or high differences between immune responses tended to be concentrated in the RBD, NTD, and HR2, the mutations with high estimated true variance were found to be most notably in the low-binding region of the RBD (outside the receptor binding motif) (Figure 3B). In this area, a mutation is less likely to be exposed to antibodies directly, but able to alter the context of the NTD and RBD, amplifying or attenuating the effects of mutations in those epitopes. Within the NTD, several deletions and substitutions at amino acid positions 24, 25, 26 had high variance; these positions are close to the NTD N-terminus (residues 14-20), and likely pivot the conformation of this known epitope.

From these results, we hypothesized that interaction may occur between two mutations with high variance, or between a mutation with high variance and a mutation with large effect size. Defining high-variance mutations as those with a corrected variance estimate greater than 15 (n=17 mutations), and high-effect mutations as the 15 mutations with greatest magnitude immune escape / capture, we described a set of 391 potentially epistatic unique mutation pairs, of which 273 were observed in the data at least 10 times. We then re-ran the Bayesian hierarchical model of immune escape including all the linear mutation terms plus these 273 quadratic terms (Figure 3C). We found the strongest immune escape interactions to occur between N764K & K444T and R408S & R346T, despite most of these linear terms having small or even negative effects on their own. The strongest immune capture interactions occurred between N969K & T376A and R408S & L452R. We compared pairwise nonlinear effects to amino acid distance and 3D distance in both open and closed RBD conformations and found that mutations could have strong nonlinear interactions regardless of their proximity (Supplemental Figure S10).

### Immune escape is one of several important factors for spreading of SARS-CoV-2

Is an increase in immune evasion enough to cause a mutation to spread through the population of circulating virus during a pandemic? We used two previously described models of mutation-specific fitness(38,39) to estimate the peak growth rate of spike mutations between the beginning of 2021 and the end of 2023. We found that the mutational growth rate estimated using Bayesian Viral Allele Selection (BVAS)(39) had a Spearman correlation of 0.25 with the Core Antigenic escape effect (p<0.05) (Supplemental Figure S11). The several mutations with low confidence in phenotype limit the magnitude of the correlation. To focus only on confidently phenotyped mutations, we select mutations with a posterior inclusion probability in the growth model of at least 0.95 and classify growth-promoting and growth-attenuating mutations based on median split of growth rate. We classify immune escape versus immune capture mutations based on the sign of the core antigenic effect, and select mutations with 95% CI core effect excluding zero. On the subset of selected mutations with either a confident growth or escape phenotype, classification accuracy of growth phenotype based on escape phenotype is 0.54. On the subset of mutations with both a confident growth and escape phenotype, classification accuracy is 0.72 (Figure 4A). Several mutations exhibited growth behavior that could be explained by immune effect, i.e. growth-promoting, immune escape mutations and growth-attenuating, immune capture mutations. Many of these mutations were in the RBD and NTD and included F486P, E484K, L452R, K444T, and R346T, which all had high immune escape and estimated peak growth, or T478Q which was considered an immune-capture mutation that was attenuative for growth.

**Figure 4:**
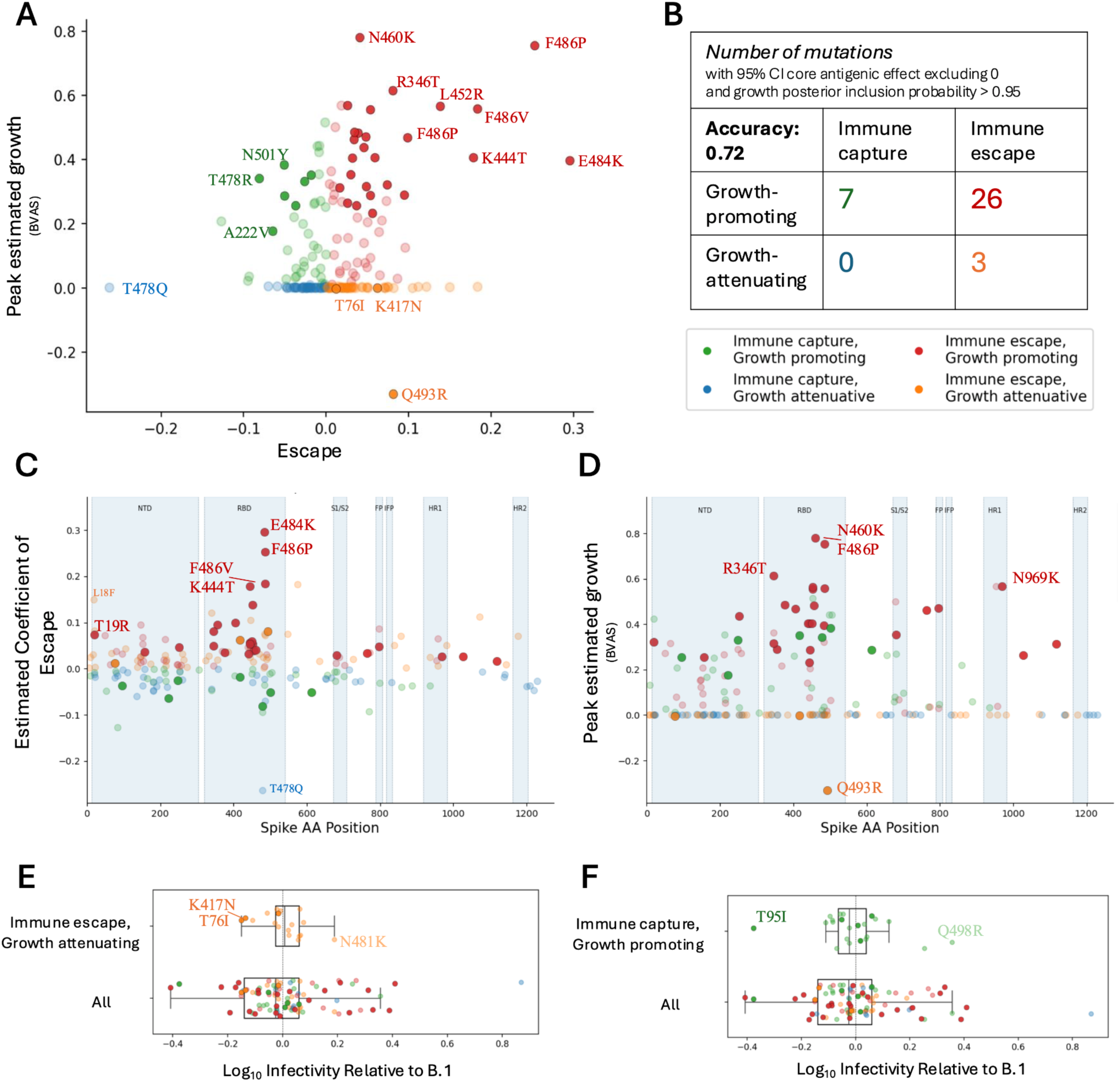
Correlation with growth estimates. (A) There is a positive relationship between estimated escape and peak growth rate of mutations. Confidently phenotyped mutations are shown as opaque. (B) Mutations can be divided into four categories based on immune escape / capture and growth. Classification accuracy of growth based on escape for confidently phenotyped mutations is 0.72. (C) In the RBD, NTD, and S2 subunit, the mutations with greatest escape are growth-promoting and those with least escape are growth-attenuating. (D) Outside of the S1/S2 cleavage site, the fastest growing mutations exhibit immune escape. (E) Some mutations, like K417N and T76I, are deleterious for infectivity, which may explain why they are growth-attenuating. (F) Mutations like T95I are deleterious for infectivity and contribute to immune capture, yet continue to promote growth.

We notice that in most regions of the spike protein, the mutations with the strongest immune effect exhibit concordant peak growth behavior (Figure 4C). E484K, F486P, and T478Q in the RBD all have growth behavior that can be explained by immune effect; K417N, T19R, and W152C all show the same pattern in the NTD; T1027I and D796H in the S2 subunit follow similarly. This trend holds for the highest growth mutations, which are generally found to be escape mutations in the RBD, NTD, HR1, and the region between NTD and RBD especially (Figure 4D). However, in the S1/S2 cleavage region, this trend reverses; the growth-promoting mutations here are all immune-capture mutations, except for A688V.

Growth estimates differed between the two models, BVAS (39) and PyR_0_ (38). BVAS, a diffusion-based variable-selection model, inherently accounts for mutational “drift” (not to be confused with evolutionary drift), the phenomenon of unfit mutations being carried along evolutionarily with more fit mutations. PyR_0_, on the other hand, is a more descriptive model that regresses counts of strains and mutations hierarchically over regions to infer growth rates. Where BVAS seeks a more explanatory coefficient of growth, indicating that a mutation may be causing growth of a strain, PyR0 yields a more descriptive metric of how much the mutation can be seen to grow in the population. Though BVAS shows few cases outside of the S1/S2 cleavage region of high-growth immune capture mutations, PyR_0_ isolates several mutations with this behavior near the NTD and RBD, including T859N, Y248N, P9L, T478R, and N703I (Supplemental Figures 12-16). This indicates that immune-capture mutations may still grow in the population if they are carried in strains that have high growth attributable to other factors.

Due to the construction of the BVAS model, mutations flagged for growth are likely drivers of growth, not passengers being carried along in otherwise fit strains. So how may we explain why some mutations that have high immune escape are still attenuative for growth? We hypothesize that some mutations may impact other phenotypes, like infectivity, which may be related to growth. To describe infectivity as a mutation-level phenotype, we built a Bayesian regression model similar to that used to infer immune escape (see Methods), and trained the model on 511 strain-level measurements of infectivity (plaque assay), consisting of 147 unique sequences spanning 213 amino acid mutations. From Figure 4E we can see that on average, mutations are deleterious for infectivity. Mutations like K417N and T76I in particular are highly deleterious for infectivity, which may prevent them from growing in the population despite the immune escape advantage they confer. On the other hand, in Figure 4F we see that there are mutations such as T95I, which are both deleterious to infectivity and immune escape, yet still contributing to growth. It is known that deletions in the NTD may serve to stabilize the rest of the spike trimer(40); perhaps the substitution T95I also provides stability, conferring a phenotypic advantage not measured in our dataset.

### As immune escape increases over time, so does growth rate of viral strains

To put these mutation-level questions into context of strain-level phenotypes, we analyzed infectivity, immune escape, and growth rate trends of World Health Organization (WHO)-defined Variants of Concern and Sublineages (VoCs) (https://www.who.int/publications/m/item/historical-working-definitions-and-primary-actions-for-sars-cov-2-variants). Among these twelve previously circulating VoCs, there are nine for which we have both (a) measurements of strain-level phenotypes, and (b) estimates of mutation-level phenotypes for all mutations in the strain. Since we don’t have adequate data on BA in our dataset, we included BA.1 in our analysis instead, leaving us ten total strains to analyze: Alpha, Beta, Gamma, Delta, BA.1, BA.2.75, BQ.1, XBB, CH.1.1, XBB.1.5. These variants are listed and analyzed in temporal order of appearance in the pandemic, where “appearance” is defined as being collected at least 50 times on a given date, as reported by GISAID(35).

Growth rate of these VoCs was found to monotonically increase with each consecutive variant (Figure 5A). A similar trend in immune escape was observed (Figure 5C), lending weight to the hypothesis that immune escape is a key driver of pandemic growth at the strain level. Immune escape, however, did not increase uniformly: some strains, such as Gamma, Delta, and XBB.1.5 had somewhat lower immune escape compared to contemporaneous variants, yet still exhibited higher growth than the variants that came before them. Though it is possible that another phenotype is responsible for the increase in growth among these variants, it seems unlikely that infectivity would be that phenotype, as there doesn’t seem to be any particular trend uniting the infectivity measurements or predictions of Gamma, Delta, and XBB.1.5 (Figure 5B). Infectivity has been observed to trend downward over time; however, the presence of high and low outliers in the earlier VoCs suggests that the trend in infectivity may be better described as “stabilizing” over time.

**Figure 5:**
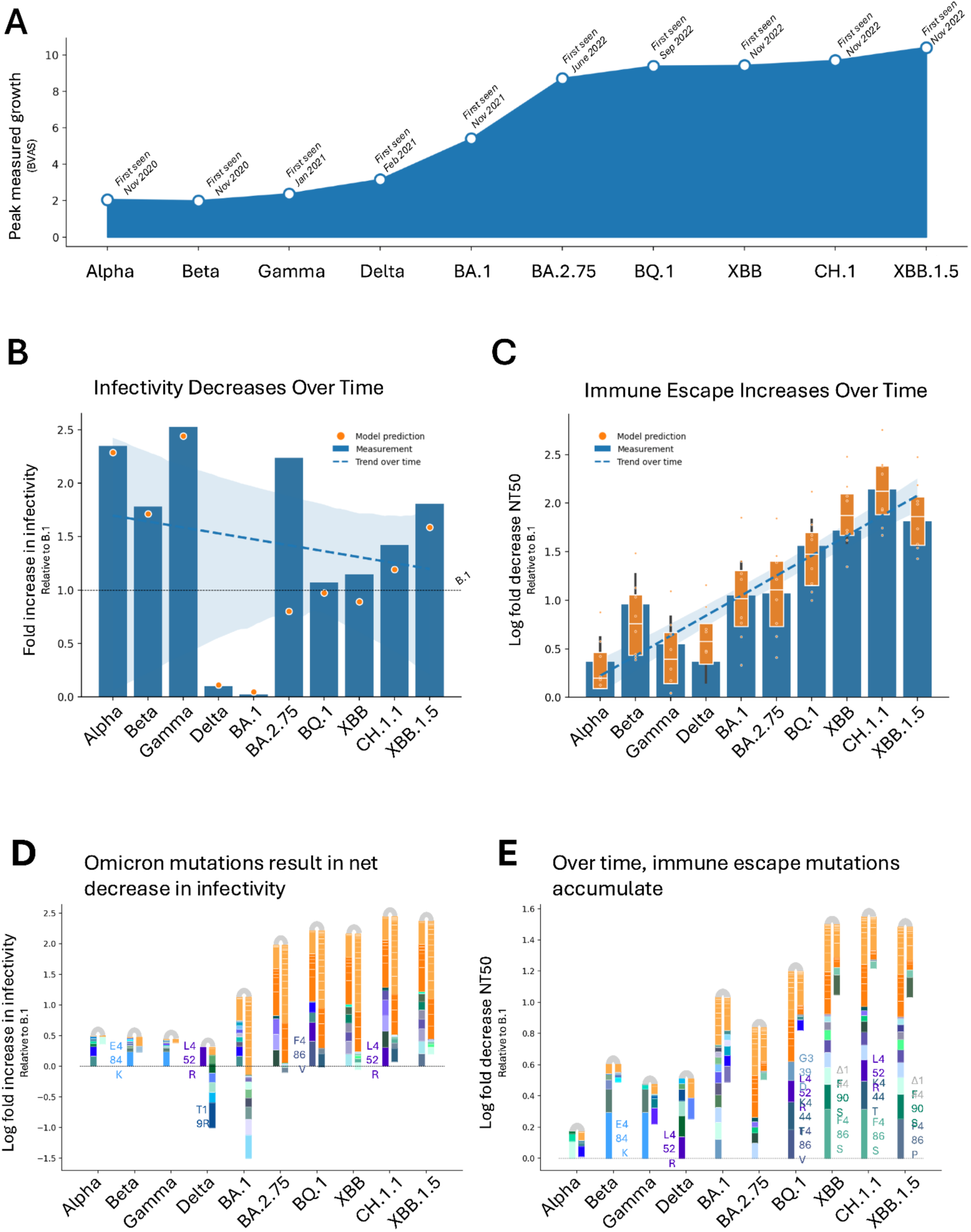
(A) Growth, as measured by BVAS (39) monotonically increases with appearance of consecutive VoCs. (B) Infectivity of VoCs stabilizes around near-neutral infectivity relative to B.1 (fold increase in infectivity relative to B.1=1) over time. Model predictions (orange dots) capture measured infectivity (blue bars). (C) Model closely predicts immune escape, which increases over time. Measurements of immune escape (log fold reduction in neutralizing titer relative to B.1, normalizing such that B.1 has log fold reduction of 0) are shown as blue bars, with range due to various serum pools, assay types, serum types, time since last exposure, and most recent variant reflected as black bars. Model predictions of immune escape are shown as orange dots, with orange boxplot summarizing differences caused by serum pool and bias terms. (D) Over time, positive infectivity mutations (left bars) and negative infectivity mutations (right bars) balance out to achieve nearly neutral infectivity relative to B.1 in later strains (log increase in infectivity = 0). This delicate balance is achieved by accumulation of several mutations – Omicron mutations which, taken as a set, decrease infectivity, and other offset mutations. Omicron mutations are shown in light orange if they are present in every VoC post-BA.1 and dark orange if they are present in every VoC post-BA.2.75. (E) While Omicron mutations (light and dark orange) eventually become fixed among the VoCs, many other immune-escape mutations are added over time to increase escape.

We ran a multivariate linear regression of growth over infectivity and immune escape and found that these variables together may account for 33% of growth of viral strains (R^2^=0.334, p<0.05). We also performed regressions of growth over immune escape or infectivity separately to better estimate the individual contributions of each phenotype, and found that when leaving out immune escape as a regressor, the R^2^ decreased by 99% (ΔR^2^= 0.3318). When leaving out infectivity, the R^2^ remained nearly unchanged (ΔR^2^= 0.0001). This further suggests that immune escape is a strong contributor to growth at the strain level.

### Mutation level analysis sheds light on constraints for mutation accumulation in SARS-CoV-2

A useful feature of our modelling approach is the ability to explain the accurate strain-level phenotypic predictions by specific mutations. This allows us to see tradeoffs at the mutation level, and indicates that the successful accumulation of mutations may be constrained by multiple phenotypes. When we examine the progression of SARS-CoV-2 mutation over time and the phenotypic changes incurred, a pattern emerges (Figure 5).

The Alpha variant was the first VoC to emerge; our analysis finds it had neutral immune escape relative to B.1 since the immune escape gains conferred by Δ144 (core antigenic effect = 0. 11) was mostly offset by the immune escape deficits caused by Δ69 (core antigenic effect = -0. 07) and N501Y (core antigenic effect = -0. 05). At the same time, the deletions Δ69-70 caused an increase in infectivity, leading Alpha to be one of the most infective early variants. However, these hallmark Alpha deletions (Δ144, Δ69-70) are absent in the Beta or Gamma variants. Those variants are instead marked by mutations E484K and L18F, which increase both immune escape (core antigenic effects = 0. 30, 0. 15) and infectivity (log_10_ fold increase in infectivity relative to B.1 = 0. 24, 0. 06). Both variants contain N501Y, slightly limiting escape. Gamma shows a similar pattern.

Emergence of the Delta variant begins to break the patterns in the Alpha, Beta, and Gamma variants. While the measured and predicted escape is on a similar or slightly lower par than the previous variants, the combination of mutations present is completely different and drastically reduces infectivity. Of the 9 amino acid mutations in Delta, only L452R increases infectivity. Even though L452R increases infectivity more strongly than any single mutation in a previously seen VoC (log_10_ fold increase in infectivity relative to B.1 = 0. 31), the other 8 mutations in Delta substantially decrease infectivity (e.g., T19R decreases infectivity by -0. 41, an even larger effect than L452R). L452R also increases immune escape (core antigenic effect = 0. 14).

BA.1, the earliest of the Omicron variants analyzed in this dataset, breaks further away from patterns observed in previous variants. This variant has the lowest infectivity, and is better at immune evasion than most previous variants. Though there are some individual mutations with large contributions to increased immune escape (G339D, Δ144, G446S, Q493R, T574K) or decreased infectivity (T95I, Δ211, Rins214EPE), most of the phenotypic shift seems to be caused by the accumulation of several mutations with small individual effects, but larger consequences in aggregate. For BA.1, there are seven mutations that offer small marginal increases in escape and decreases in infectivity: N440K, S477N, N679K, N764K, D796Y, Q954H, and N969K; yet taken together, these seven mutations lead to an increase in immune escape of 0.23 and a decrease in infectivity of -0.49. BA.1 has notably low infectivity.

This marks a new paradigm in SARS-CoV-2 evolution. Starting at BA.1, the number of mutations observed in each VoC is quadruple what was previously counted. While the 18 mutations shared in each Omicron subvariant (light orange mutations in Figure 5) cause a large decrease in infectivity (aggregate infectivity effect = -0. 89) and offer nearly no improvement to immune escape (aggregate escape effect = 0. 09), many of the remaining mutations increase both infectivity and escape.

After the emergence of BA.1, ten more mutations reach fixation in the Omicron sublineages BA.2.75, BQ.1, XBB, CH.1.1, and XBB.1.5 (dark orange mutations in Figure 5). These additional mutations increase immune escape and decrease burden of noninfectivity; as a whole, the now 28 mutations that are constant in the last five VoCs present an aggregate infectivity effect of -0. 78 and an aggregate escape effect of 0. 30. Additionally, each Omicron VoC sublineage features a quiver of mutations with strong individual contributions to immune escape and infectivity.

Despite the modest infectivity or immune escape effects of these mutations marginally, they do exhibit high growth rates. The mutations that grew to fixation in later omicron sublineages have higher growth rates on average compared to all other mutations, and they are also much more likely to be important in the growth rate prediction (posterior inclusion probability closer to 1) (Supplemental Figure S17).

## DISCUSSION

In summary, we have systematically modeled the immune escape, infectivity, and growth of SARS-CoV-2 by identifying mutation-specific influences on these phenotypes using GISAID(35) and public datasets of thousands of mutations and hundreds of escape and infectivity measurements. By employing a hierarchical model of serum pools representative of various SARS-CoV-2 immune histories, we infer serum pool-specific and core antigenic effects (marginal effects) that reveal the molecular and structural determinants of immune escape.

While we find mutations in the RBD and NTD most strongly modulate escape, aligning with prior evidence (18,29,32,41,42), we also find clear evidence for antibody activity outside these regions, including in the stalk. The HR2 region was found to impact immune escape, perhaps due to its critical role in adjusting protein conformation (43), and has been previously suggested as a useful target in nanoparticle vaccine design (44). We further investigated serum pool effects and found similarities between the immune responses elicited by Alpha, Beta, and BA.2.12.1 exposures, while Delta exposure elicited a more differentiated effect.

Overall, our estimates suggest 33.4% of strain-level variation in growth is caused by infectivity and immune escape. The remaining two thirds of growth may be explained by other viral phenotypes and by human behavior, such as travel patterns, vaccination rollouts, and other public health interventions. Variance decomposition analysis revealed that the mutations between the NTD and RBD had the greatest variance in their core antigenic effects after accounting for collinearity and lack of data. Strong nonlinear interactions between distant pairs of mutations suggest that mutations in the NTD and HR1 may stabilize the protein or alter conformation to amplify or reverse the effect of a mutation in the RBD, as has been shown to occur with D614G (45). It is important to note that all strains tested in this investigation contain the mutation D614G and that mutations are defined relative to B.1 (S:D614G), so effects described here are already in the context of any stabilization or amplification caused by D614G.

Our modeling approaches, which have included hierarchical handling of immune histories as well as variance decomposition analysis and quadratic modeling, have demonstrated that a well-designed model can reveal relevant biological features. Certain mutations in the NTD and RBD appear to be epistatic and influence the effect of other mutations. Although these effects are relatively minor, they highlight networked residues with allosteric effects. While additional model parameters can improve fit and describe biology, a simple model is well-powered to describe the general trends in immune escape, highlighting that the progressive accumulation of small, additive effects in Spike is the dominant force driving escape and growth, a finding which aligns with previous research (38,39). While more complex models can be useful for biology, they may not be strictly necessary for predictive or descriptive tasks. These findings argue for including simple, interpretable linear regression models in viral surveillance efforts.

Strengths of our approach include quantitative, systematic, and principled estimates of immune escape. Another advantage is that mutation-specific effects are readily interpretable, highlighting areas of Spike that are concordant with models of neutralization derived from structural and biological data. The approach is readily generalizable to other viruses and experimental systems. Compared to conventional regression models, our Bayesian approach also naturally accommodates a hierarchy over immune serum pools to capture immune history-dependent effects and readily quantifies the uncertainty in these effects. While typically viewed as a nuisance parameter, we show that the variance of mutation-specific effects is a useful measure and captures different biology compared to other parameters. Variance decomposition allowed us to estimate the true variance in the biological signal versus variance caused by collinearity or low prevalence of data, highlighting potentially epistatic mutations between the NTD and RBD. This was made possible by the reliable variance estimates from MCMC traces; we compared the results to those yielded by stochastic variational inference (SVI) and found the SVI results to underestimate variance, a known problem. Important effects were lost with SVI, suggesting that methods that exactly model the posterior such as MCMC may be worth the tradeoff of additional computational resources to elucidate biological effects.

Our findings contextualize broader dynamics of the pandemic during what appears to be a transition to endemicity. Mutational infectivity of VoCs increased early in the pandemic but has plateaued over time, particularly marked by the paradigm shift beginning with BA.1 where the total number of mutations balloons. Omicron sublineages had a large number of mutations with a largely zero-sum infectivity effect, suggesting that infectivity may have reached a maximum with additional mutational gains offset by mutational losses in infectivity. Infectivity has reached a local optimum whereas gains in immune escape have continued without yet reaching a bound. Transmission, therefore, has effectively ascended along an evolutionary ridge in the fitness landscape defined by escape and infectivity, undergone a “phase change,” with early variants benefitting from the combined effects of enhanced infectivity and escape while growth of later variants in the population is purely a result of escape. It is notable that fitness gains in escape have not come at the cost of infectivity even though Spike is both the ligand for binding to the viral entry receptor and the immundominant antigen; this suggests that SARS-CoV-2 will be able to participate in a prolonged cat-and-mouse game of immune escape without incurring costs in infectivity, a dynamic which is likely to facilitate endemic circulation of the virus in the human population.

## METHODS

### Data sources

We collected data on two phenotypes: infectivity and immune escape. We used infectivity and immune escape data collected by Youssef et. al., 2024(5), wherein several combinations of mutations to the SARS-CoV-2 spike protein were computationally designed and tested *in vitro* for infectivity and immune escape using previously described assays(5,45–48). Serum used for the immune escape neutralization assays was collected from the POSITIVES study(49,50). We also used immune escape data made available by the Stanford University Coronavirus Antiviral & Resistance Database (Cov-DB)(33) which collates previously published experimental immune escape data across various strains of SARS-CoV-2. Across both this data source and the neutralization assays performed by Youssef et. al., 2024(5), we only used the subset of experimental data available where exact fold reduction in neutralizing titer was known (‘Fold Reduction: Cmp’ was ‘=’), exact mutations were known, assay control for calculating fold reduction was wildtype or B.1 (wildtype with D614G), host was human, and there was at least one mutation present. For all viral strains analyzed, the closest Variant of Concern / Base VoC was previously reported. Regarding serum used for neutralization assays, the Stanford Cov-DB(33) provided the number of previous exposures to SARS-CoV-2, and for the POSITIVES serum donors, number of exposures was extrapolated from immune history variables such that donors with breakthrough infections or booster vaccines were assumed to have had three exposures, vaccinated donors were assumed to have had two exposures, and donors from whom convalescent plasma was collected were assumed to have had one exposure. Months since most recent exposure was provided by the Stanford Cov-DB, and was assumed to be 2-6 months for the POSITIVES participants. Assay type was provided by the Stanford Cov-DB, and assay type is known to be Pseudovirus (lentivirus) for all results from Youssef et. al., 2024. We also used metadata associated with 17,243,486 sequences available on GISAID up to August 8, 2025, via gisaid.org/EPI_SET_250808ux.

### Bayesian hierarchical regression model of immune escape

We regressed escape as a linear function of genetic changes in the Spike protein. Mutations correspond to covariates in the model. We used a Bayesian model with Laplace priors over mutations to achieve regression shrinkage and variable selection. We implemented the model hierarchical over pools that account for immune histories by defining pool-specific (serum-level) and pool-agnostic (core antigenic) effects. The model was implemented in the python probabilistic programming language pyro(51). Hyperparameters were set by examining the conditional posterior distribution across a range of hyperparameters and by optimizing parameter stability. Each mutation was modeled with a Core Antigenic Effect describing the mutation’s average immune escape across all antibodies (Equation 5), and a Serum Pool Effect describing the mutation’s immune escape from antibodies elicited by an individual’s immune history (Equation 4). Intercept terms were included in the model to adjust for noise in the experiments (such as different assay types) and noise in immune histories of plasma / serum donors, such as previous viral exposure via vaccine versus infection, number of previous exposures, and months since last exposure.

Specifically, we estimated log fold reduction in neutralizing titer in serum pool *j* as the sum of mutation serum pool *j* effects plus intercept terms using

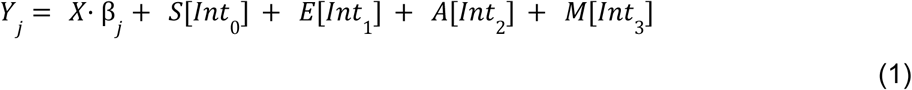

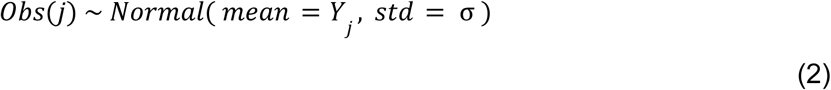

so that observations are sampled from a normal distribution with spread

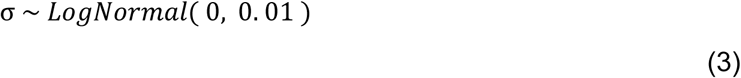

and observations are centered on the predicted escape value *Y*_*j*_ ∈ ℝ of all *N=5,223* constructs from serum pool *j*. The matrix *X* ∈ {0,1}^*N*⨯*F*^ is the design matrix specifying presence or absence of the *F=779* mutations, and β_*j*_ ∈ ℝ^*F*^ hold the serum pool effects for pool *j* of all *F* mutations. *Int* is a 4 ⨉ *N* matrix with each column containing the indices of corresponding intercept terms to be used in each experiment.

Intercept terms are encoded as vectors of intercept values *S* ∈ ℝ^3^, *E* ∈ ℝ^2^, *A* ∈ ℝ^10^, and *M* ∈ ℝ^3^, each sampled from a standard normal distribution of appropriate sizes, to reflect the three possible serum types (unknown, vaccine, convalescent), two possible exposure types (<=2, >2), 10 possible assay types (pseudovirus, pseudovirus (lentivirus), pseudovirus (HIV), pseudovirus (MLV), pseudovirus (VSV), VSV chimeric virus, hiVNT, virus isolate, VLP, SARS-VoC-2 recombinant), and the 3 possible number of months since most recent exposure (1 mo, 2-6mos, >6mos).

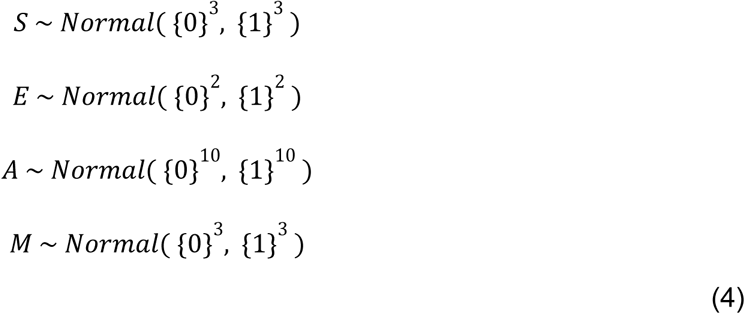

These values are shown in Supplemental Figure S2. The matrix *Int* holds the index values that describe the intercept terms which should be considered for each experiment, such that

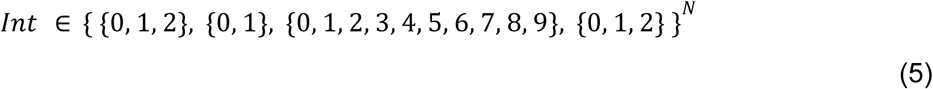

Serum pool effects are sampled from core antigenic effects, using a Laplace distribution parameterized by an inferred scale parameter:

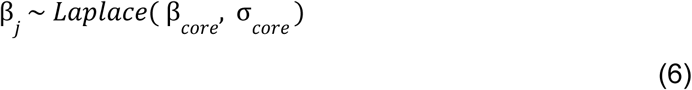

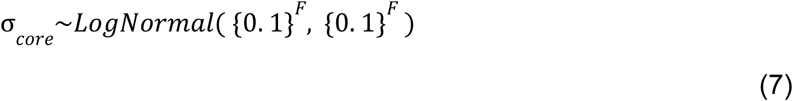

and core antigenic effects are sampled from an uninformative Laplace shrinkage prior:

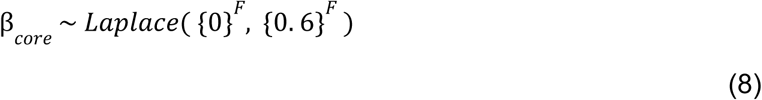

The scale value of 0.6 was determined by parameter sweep.

### Bayesian hierarchical regression model of infectivity

Analogous to the escape model, we regressed infectivity as a linear function of the sum of mutations present in a given lineage. We used a Bayesian model with Laplace priors over mutations to achieve regression shrinkage and variable selection. The model was implemented in pyro(51). It is specifically expressed as:

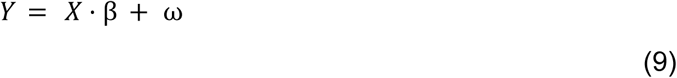

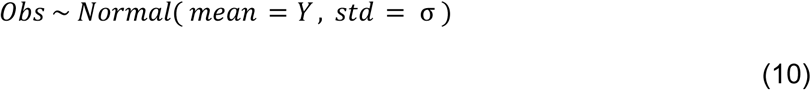

where the observations *Obs* take on the form

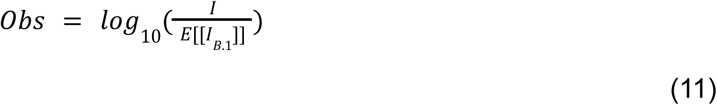

In eq. (11), *I* is the measured infectivity of a strain, and *E*[[*I*_*B*.1_]] is the average infectivity of B.1, thus observations are the log of the relative infectivity. This construction means that mutation effects β act multiplicatively on infectivity in linear space and infectivity is always positive, and we anticipate that the intercept term ω should be close to zero. Explicitly,

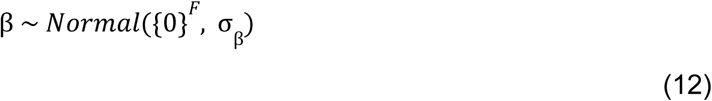

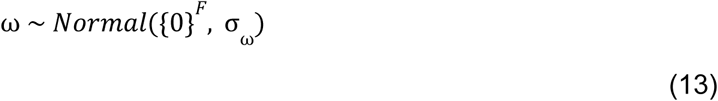

We sample all of our standard deviation terms from non-informative, uniform, positive priors:

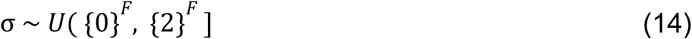

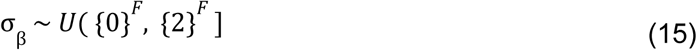

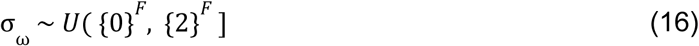

### Bayesian models of growth

We ran previously described Bayesian growth models(38,39) at multiple timepoints (approximately biweekly) throughout the pandemic. Since growth rate estimates of mutations or strains vary over time, we summarized mutation and strain growth as the maximum regressed estimate.

## ACKNOWLEDGEMENTS

The authors would like to thank members of the Luban, Seaman, Marks, and Lemieux labs. We would also like to acknowledge the curators of the Stanford Coronavirus Resistance Database and the scientists whose data is made public there. We gratefully acknowledge all data contributors, i.e., the Authors and their Originating laboratories responsible for obtaining the specimens, and the Submitting laboratories for generating the genetic sequence data and metadata and sharing via the GISAID Initiative, on which this research is based.

## Supplemental Tables

**Table S1:Serum Pool and Core Antigenic Effects for each mutation**

**Table S2: Variance decomposition of each mutation**

**Table S3: EPI_SET**

## AUTHOR CONTRIBUTIONS

Lemieux, Marks, Luban, and Seaman jointly conceptualized the project. Kotzen developed the models, performed all analyses, and wrote the manuscript. Youssef and Gurev contributed to design of analyses and model development discussions and provided feedback on statistical and computational methods. Jaimes, Luban, and Seaman provided biological interpretation and contextual guidance while Lemieux and Marks provided feedback and guidance on Bayesian modeling and analysis methods. All authors discussed the results and contributed to manuscript review and editing.

