## Supplementary figures and images for "Transition from infectivity and immune escape to pure escape as an evolutionary strategy during the COVID-19 pandemic"

### DMS-Escape.png

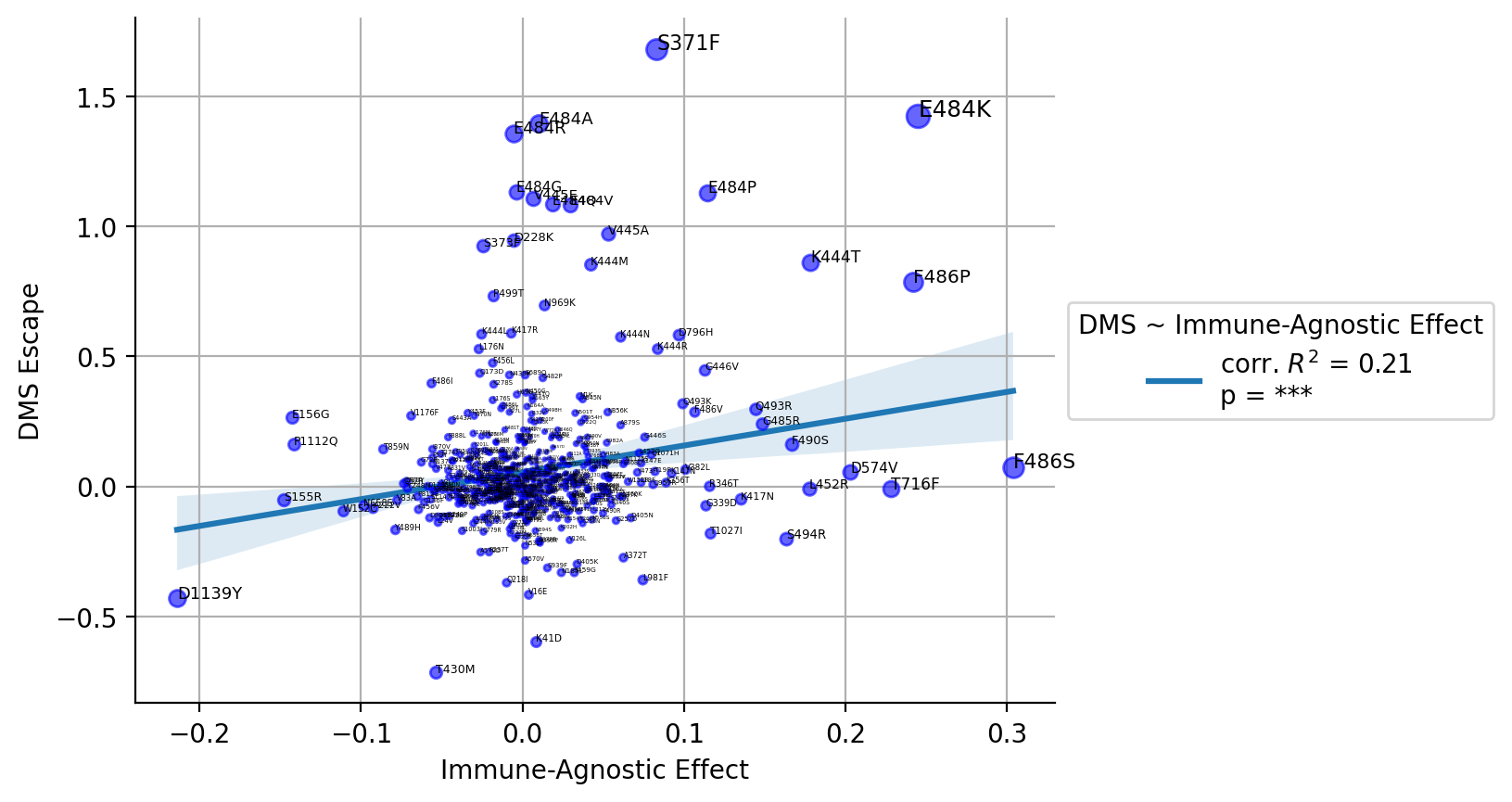

### PyR0-DMS.png

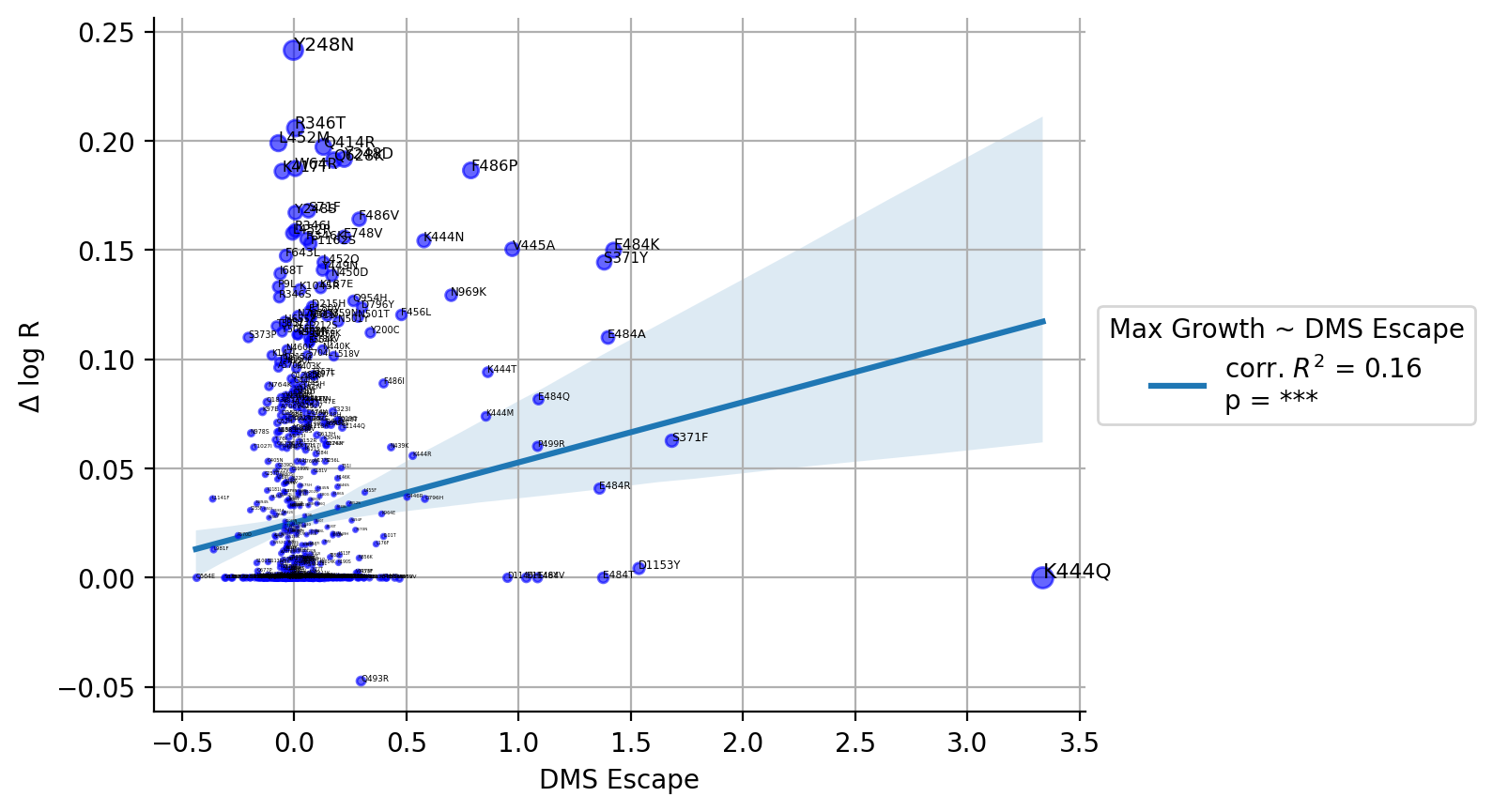

### PyR0-Escape.png

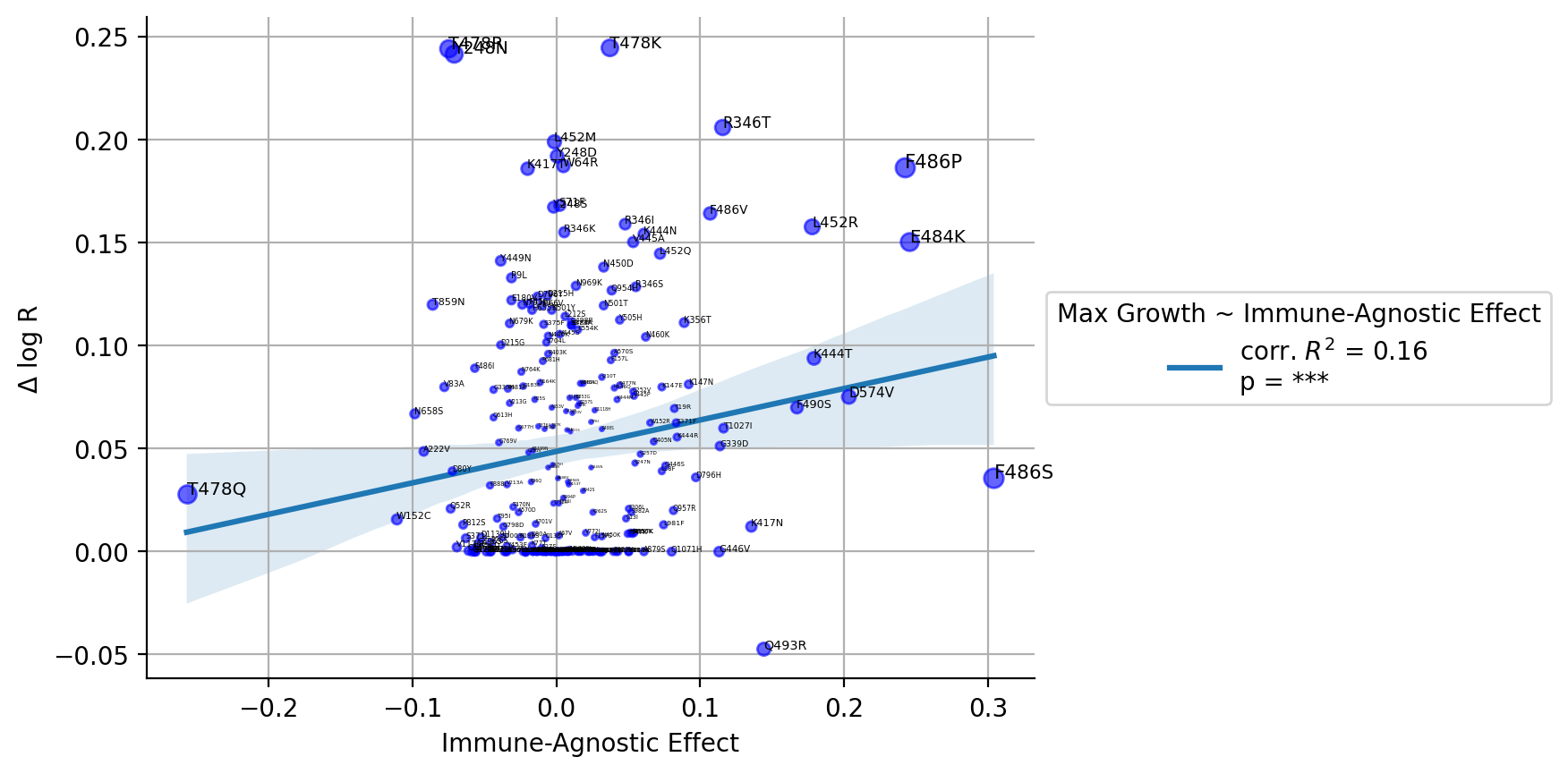

### S1.png

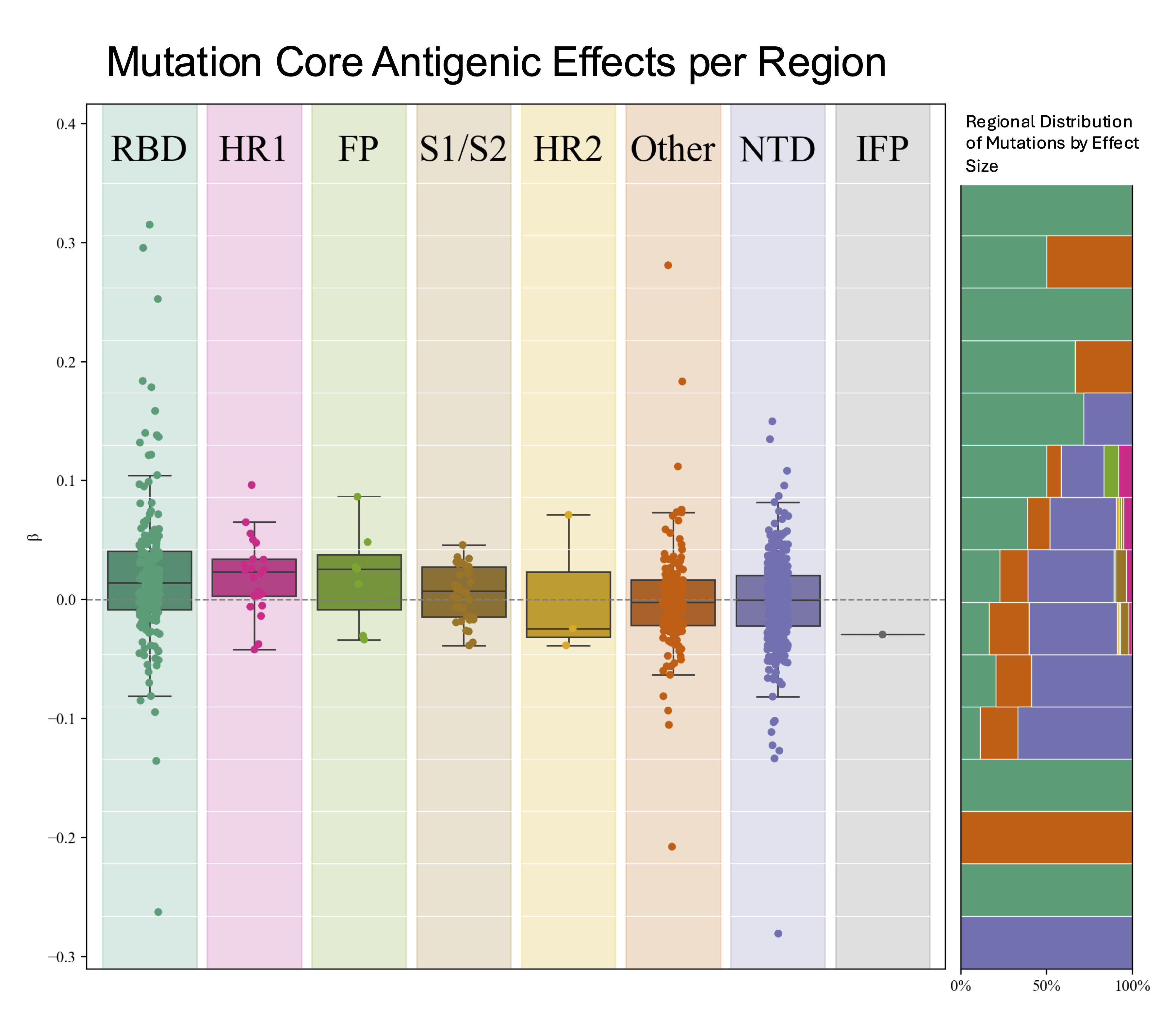

### S2.png

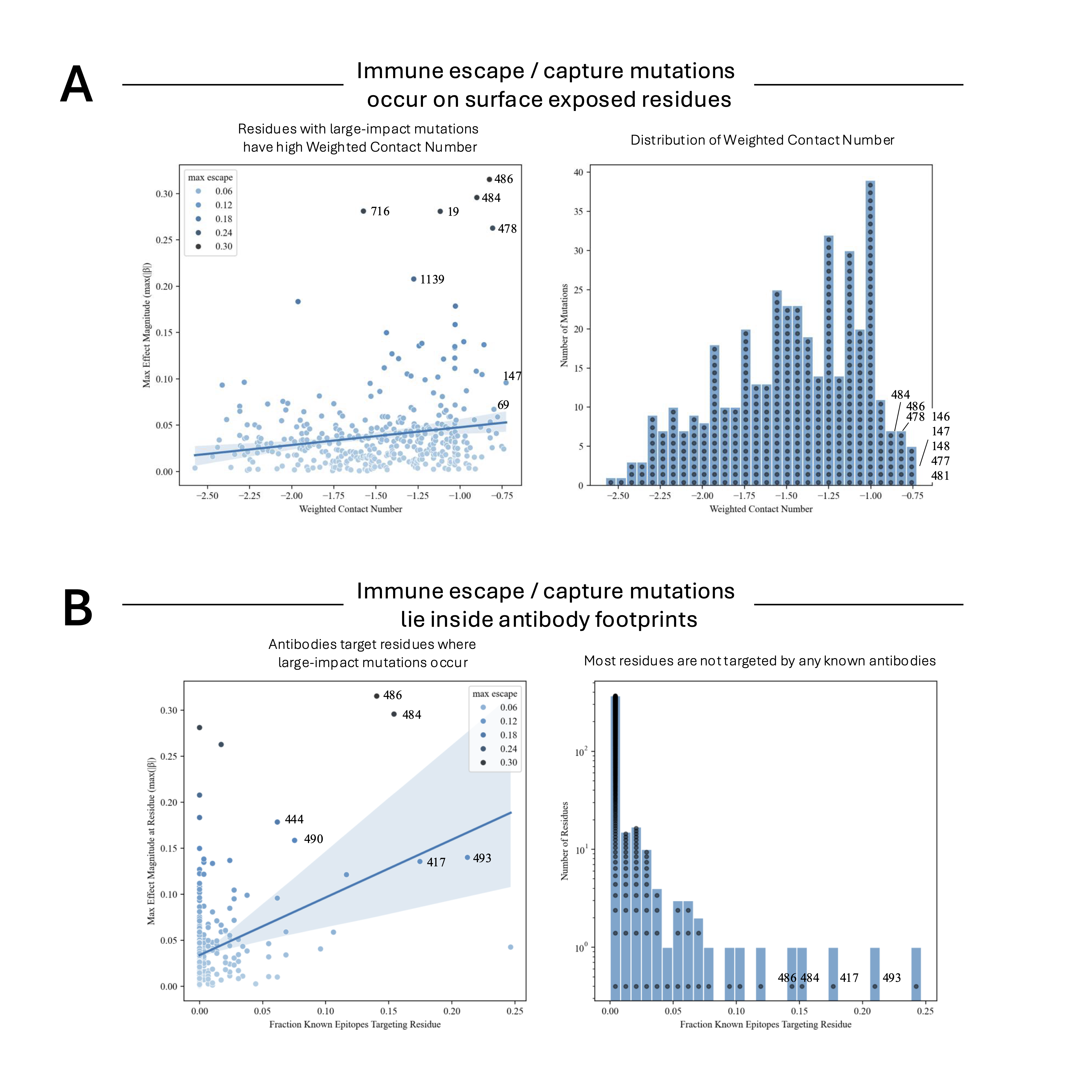

### S3.png

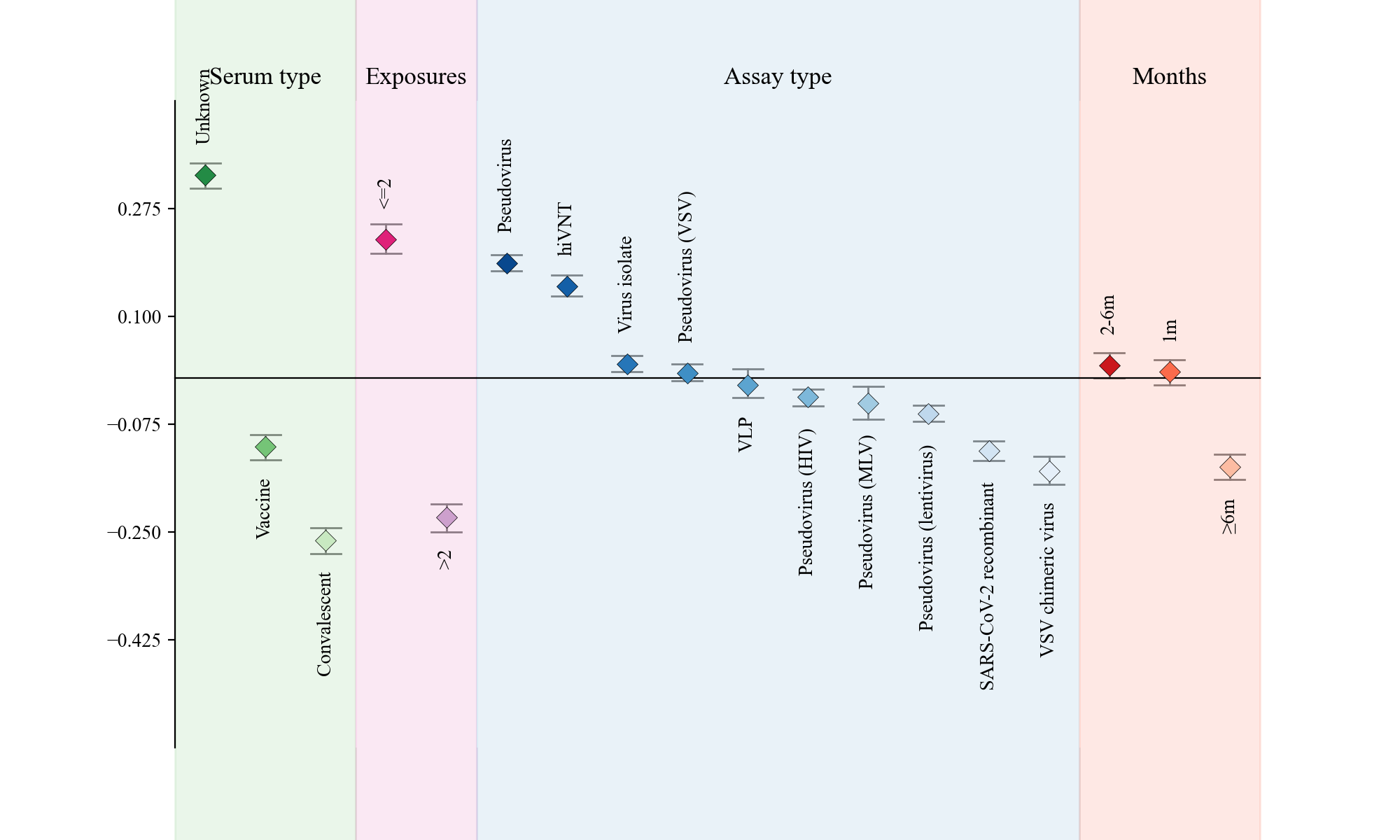

### S4.png

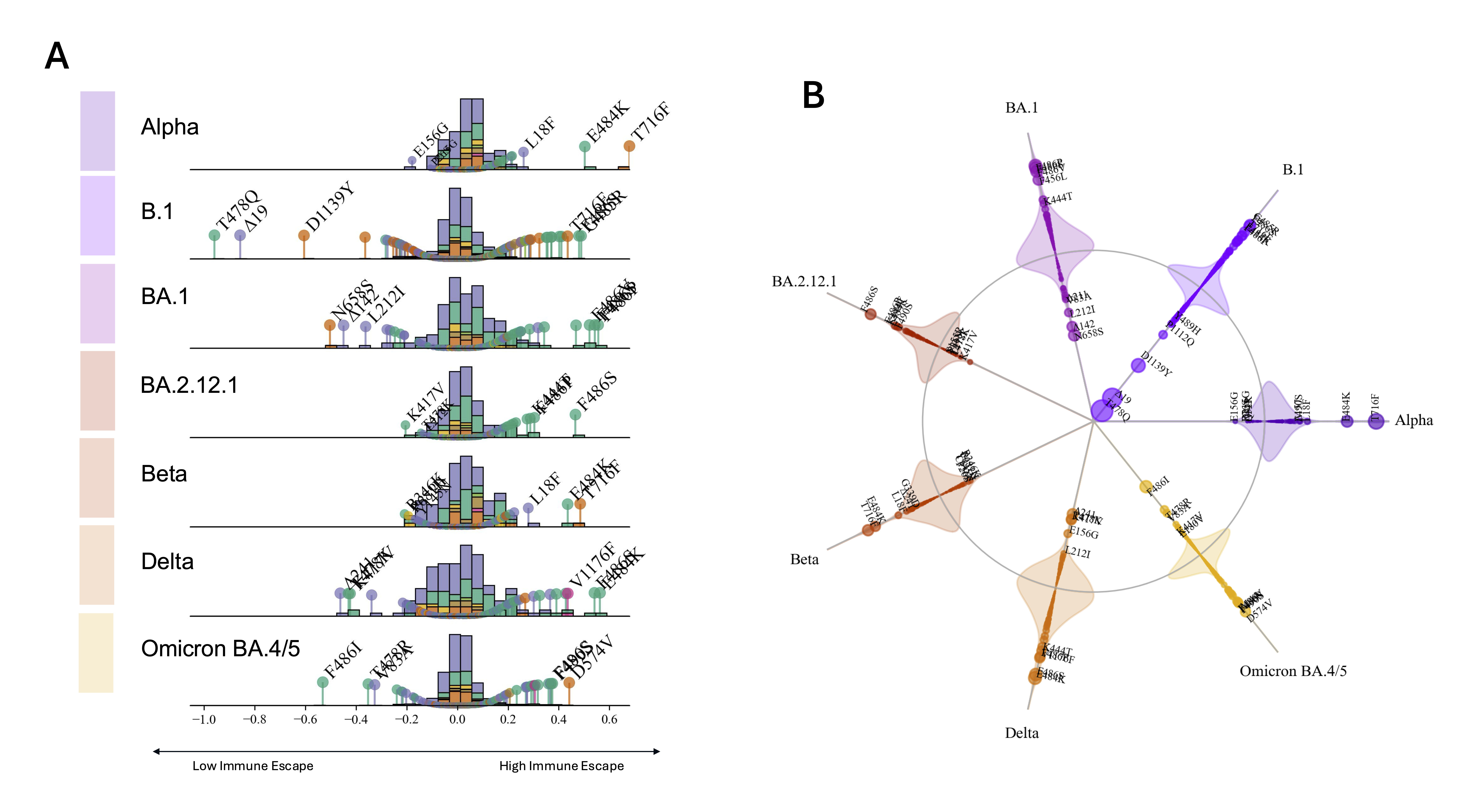

### S5.png

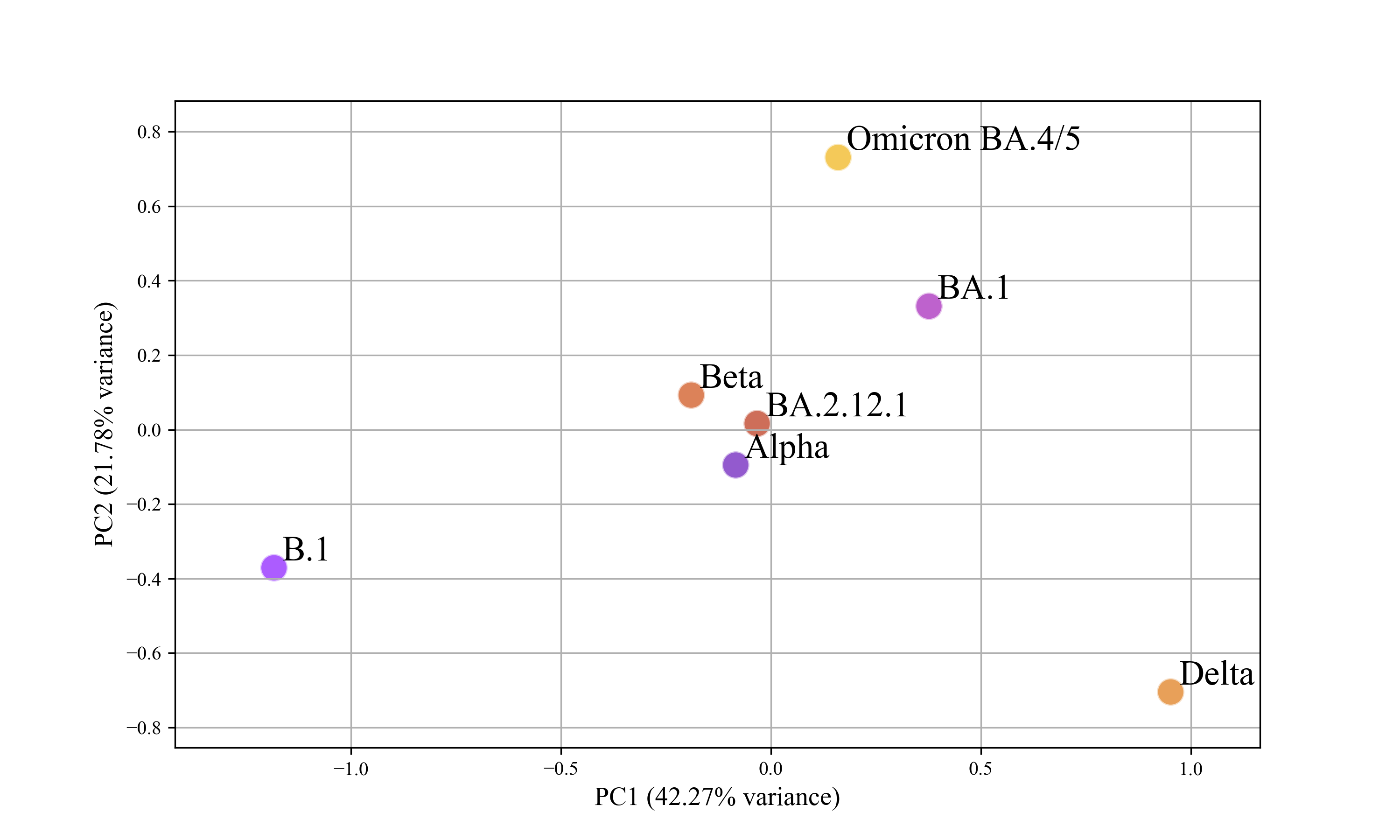

### S6.png

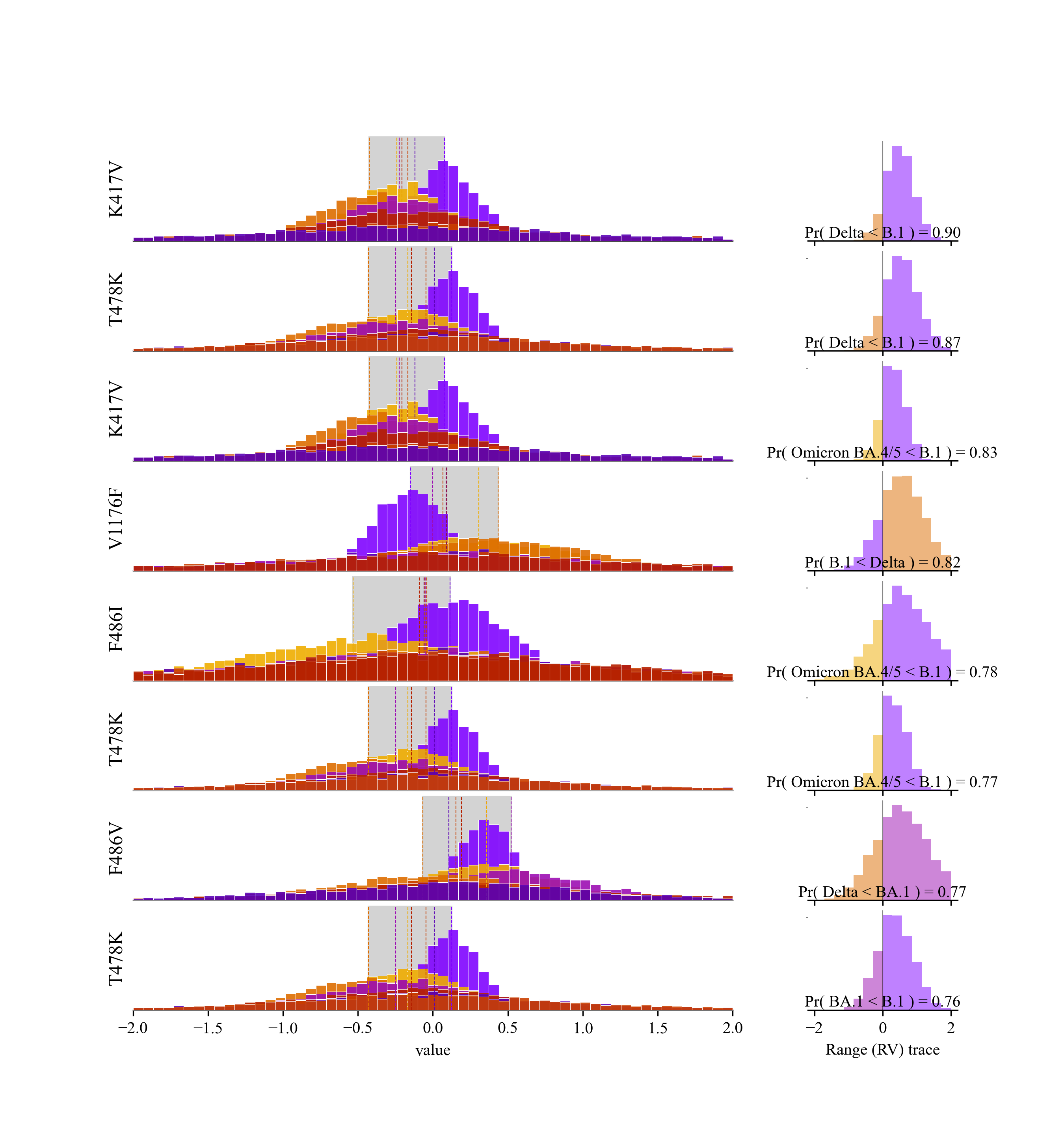

### S7.png

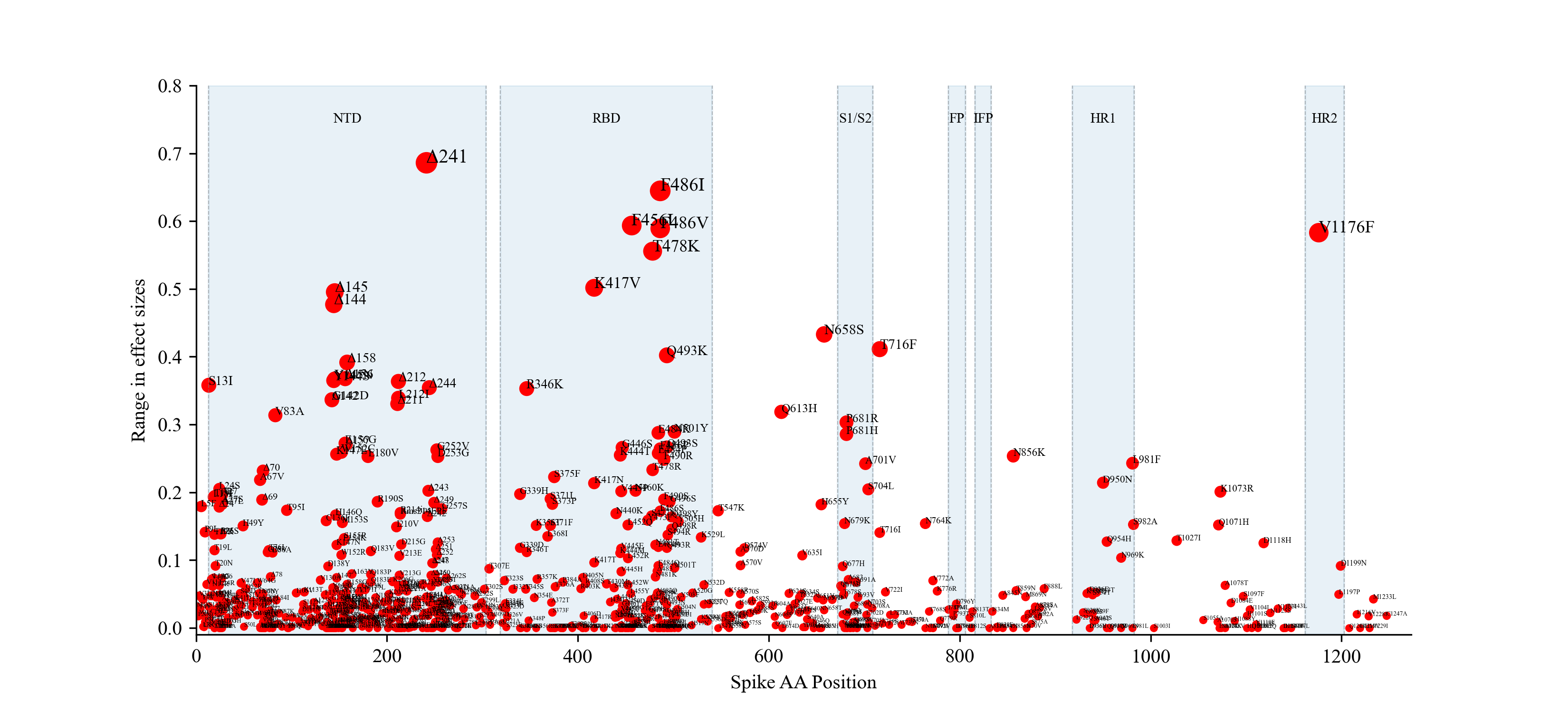

### S8.png

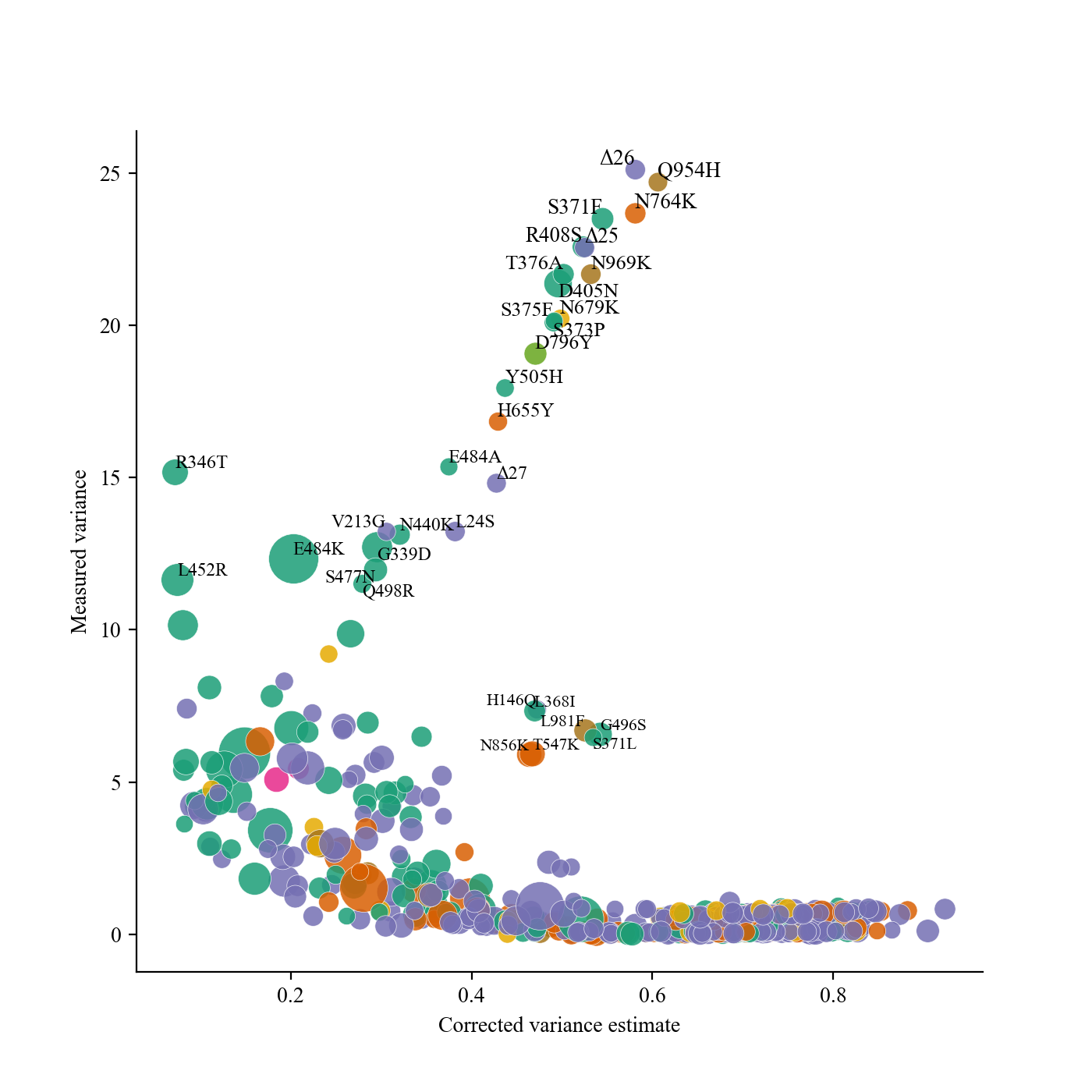

### S9.png

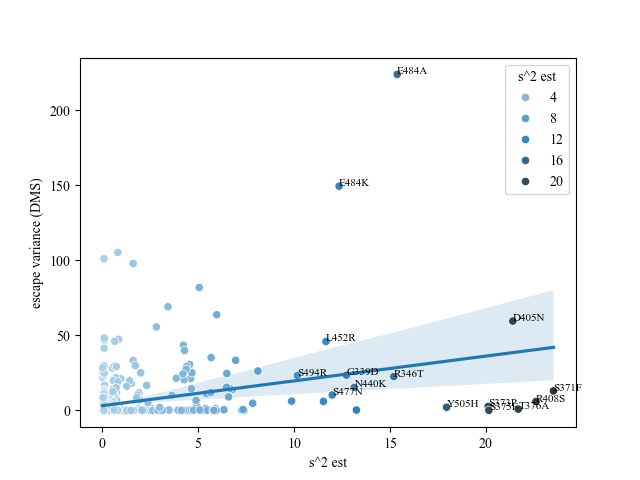

### S10.png

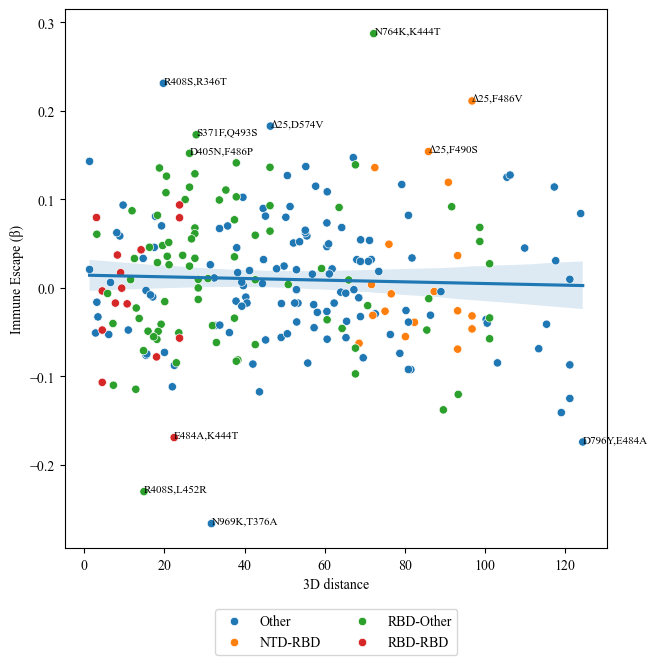

### S11.png

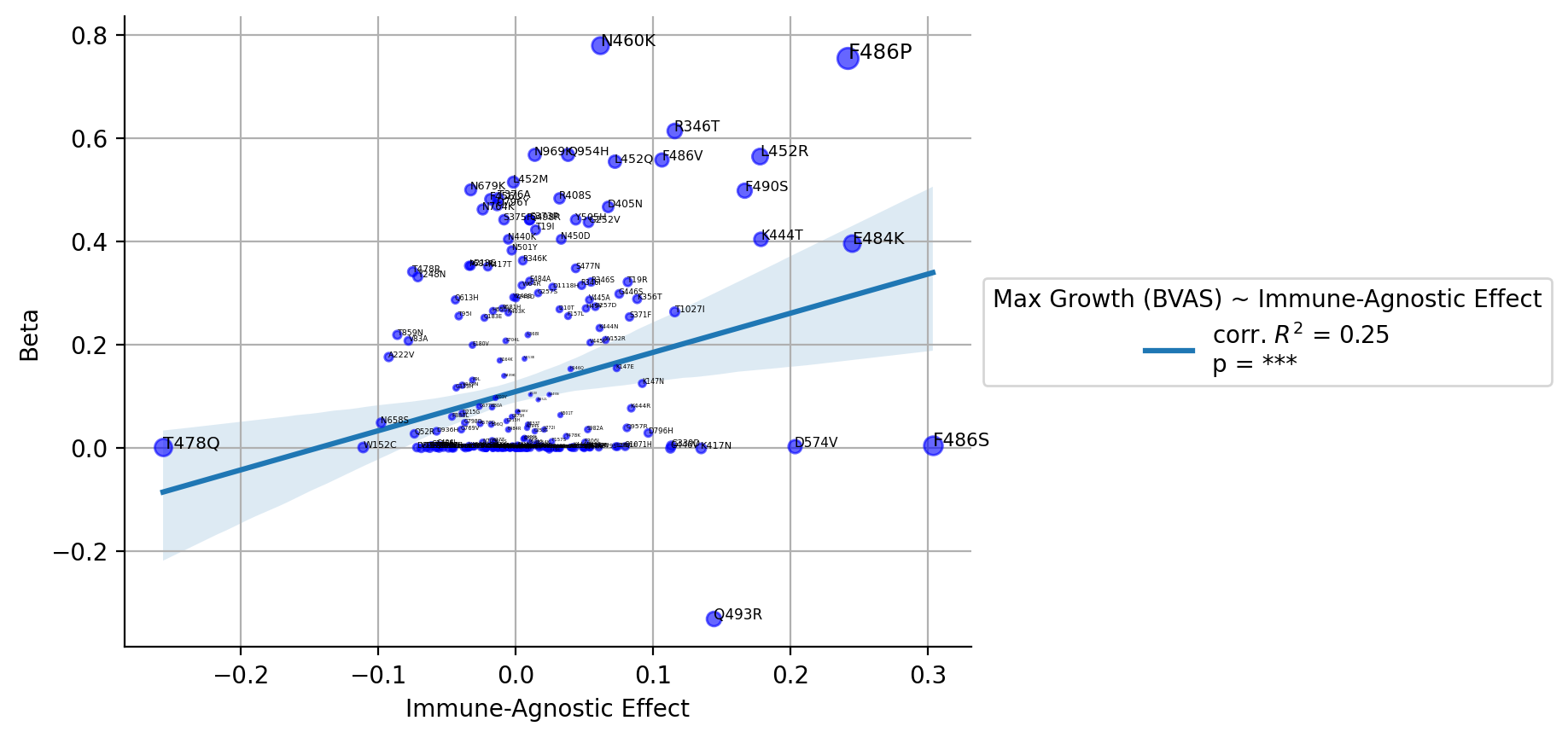

### S12.png

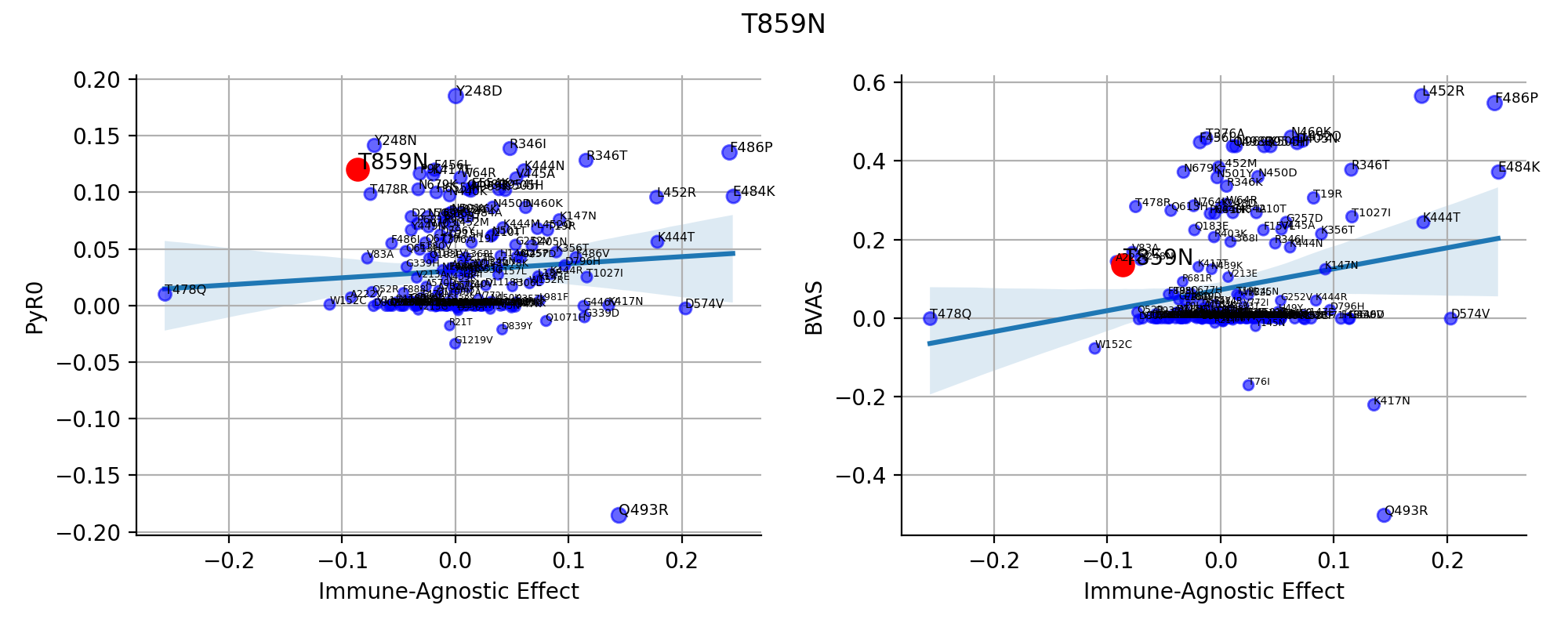

### S13.png

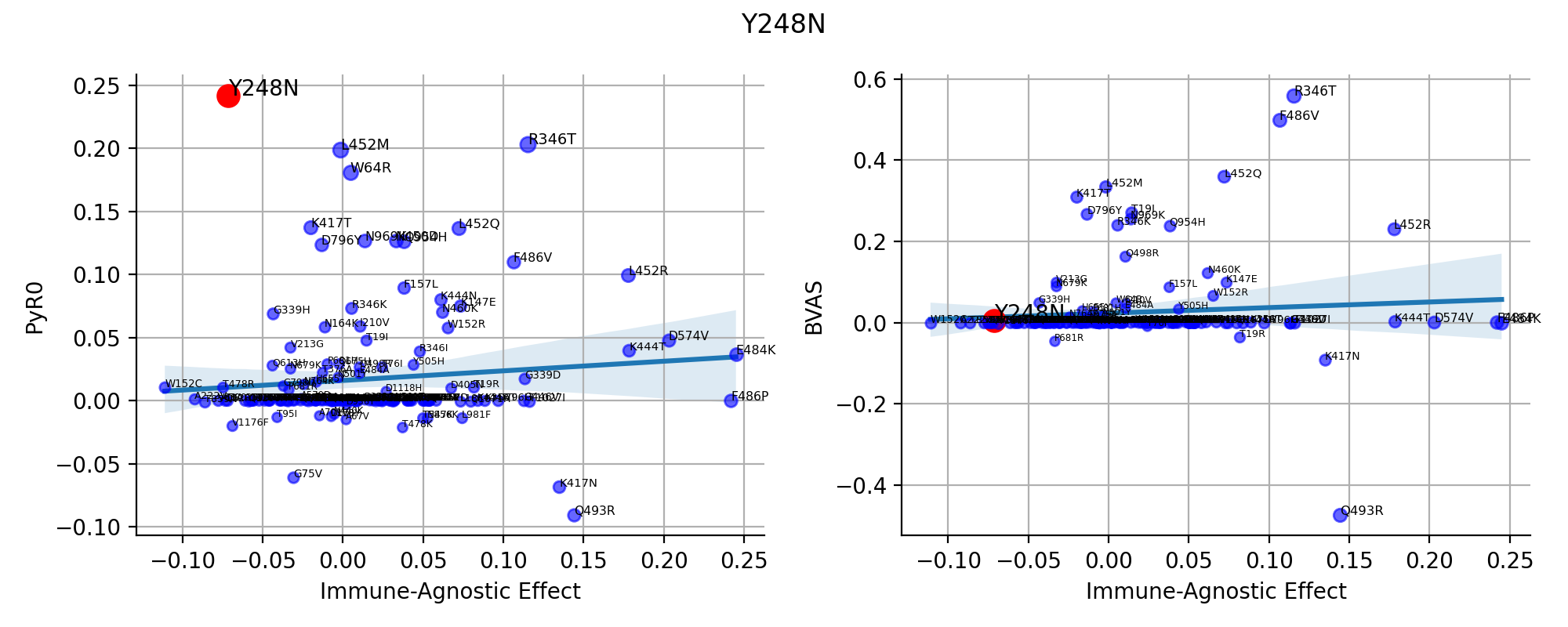

### S14.png

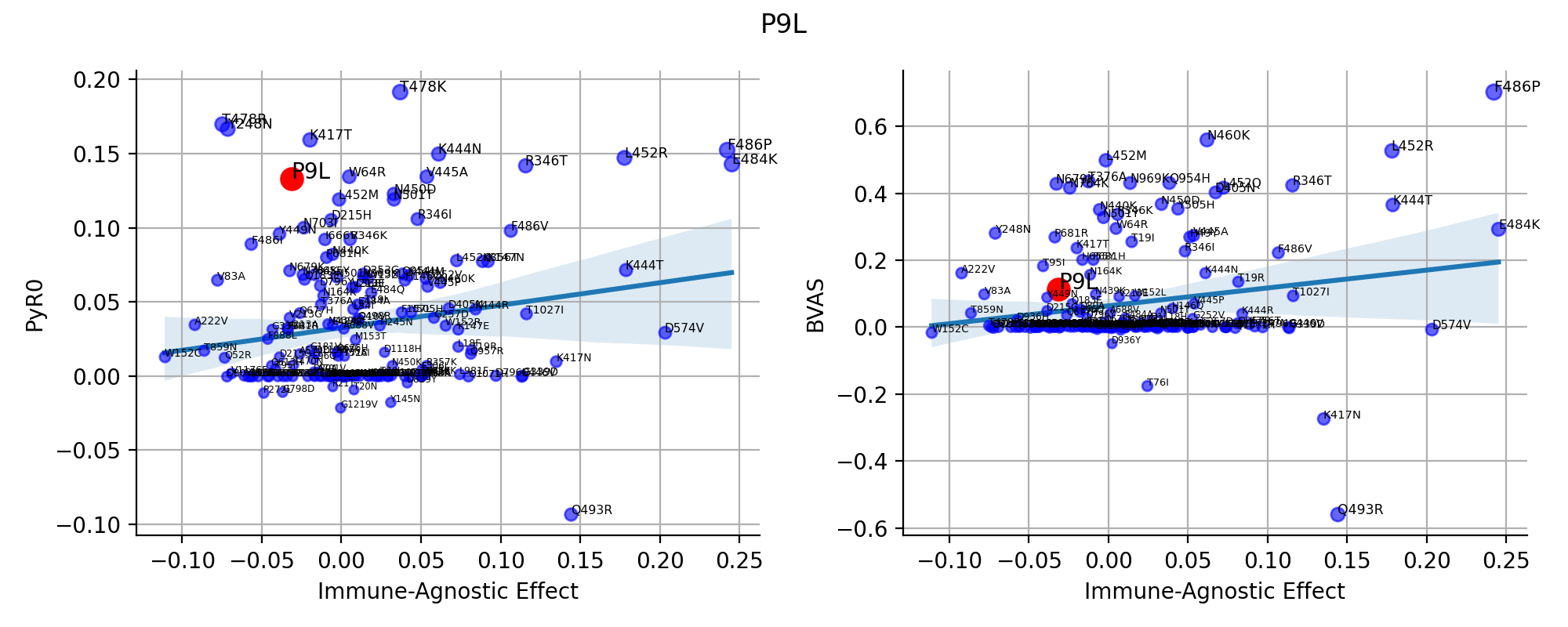

### S15.png

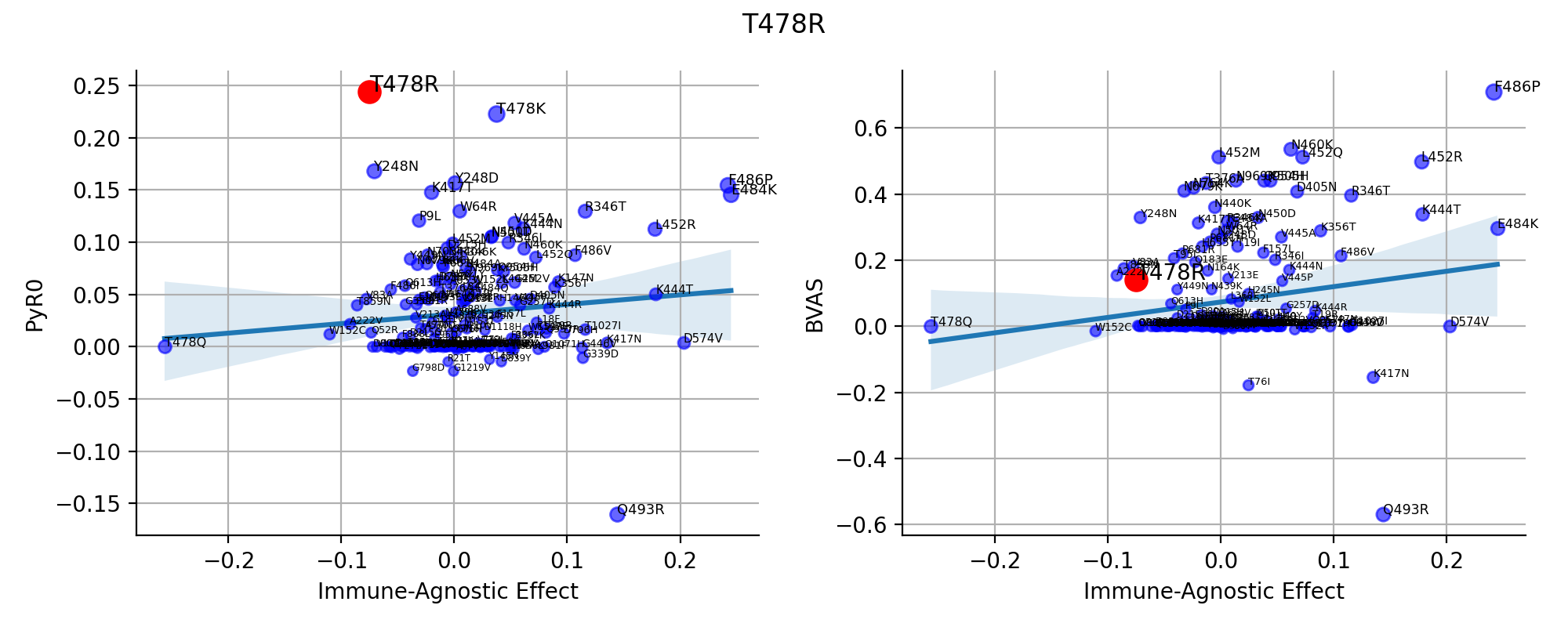

### S16.png

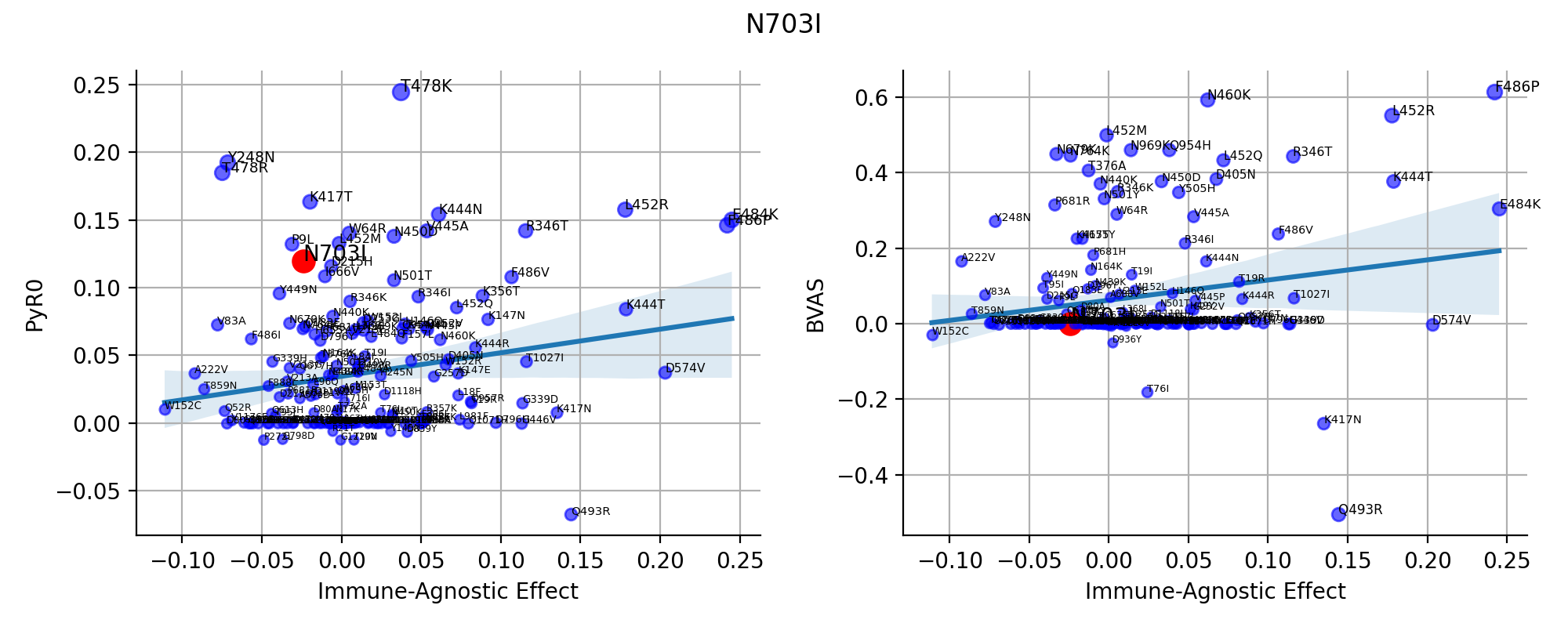

### S17.png

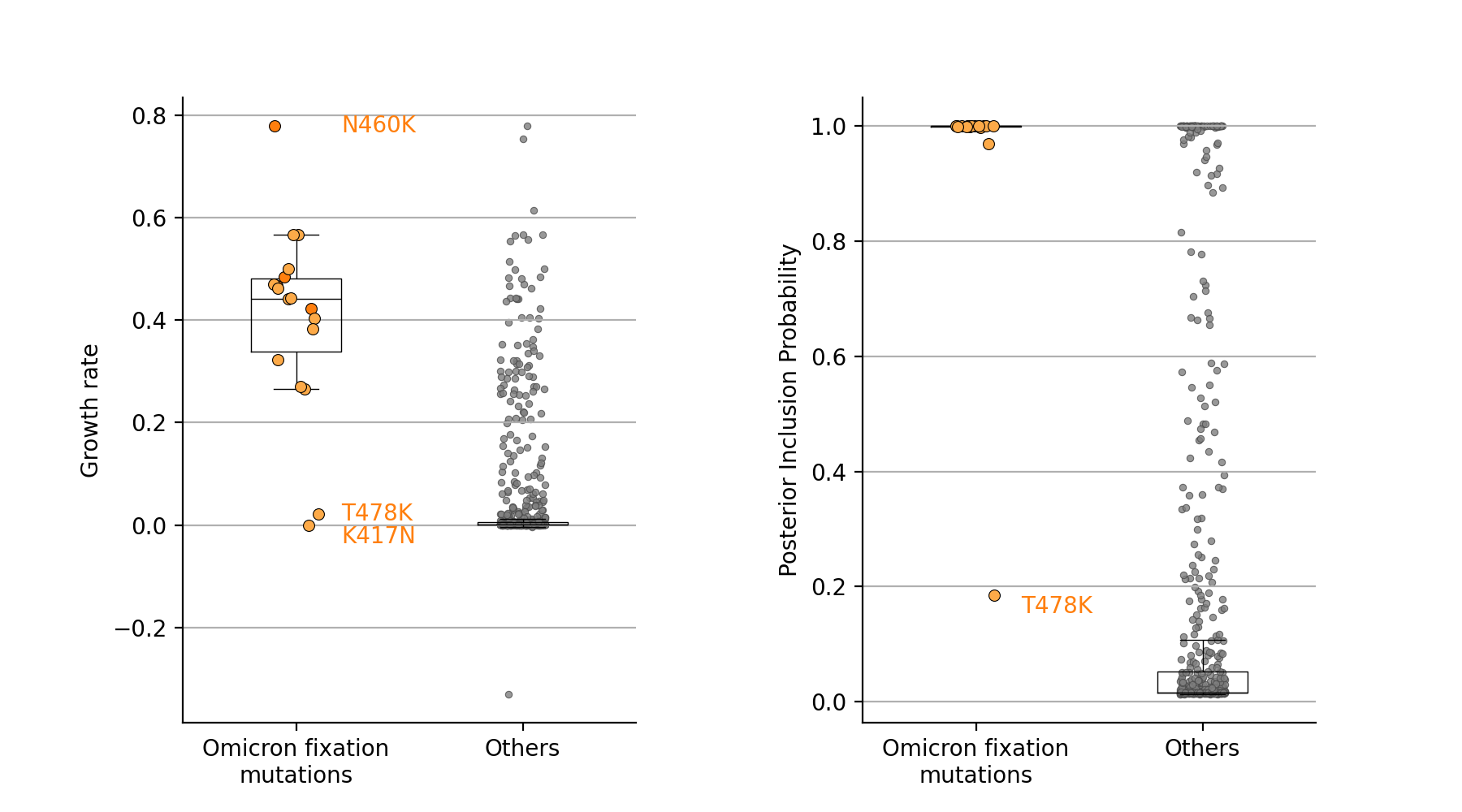
