## Supplemental Table for "Transition from infectivity and immune escape to pure escape as an evolutionary strategy during the COVID-19 pandemic": S3.pdf

### Data Availability

GISAID Identifier: EPI_SET_250808ux
DOI: https://doi.org/10.55876/gis8.250808ux

All genome sequences and associated metadata in this dataset are published in GISAID's EpiCoV database. To view the contributors of each individual sequence with details such as accession number, Virus name, Collection date, Originating Lab and Submitting Lab and the list of Authors, visit EPI_SET_250808ux

### Data Snapshot

EPI_SET_250808ux is composed of 17,243,458 individual genome sequences.
The collection dates range from 2019-12-05 to 2025-06-15;
Data were collected in 222 countries and territories.
